# Evolutionarily Conserved Fragments reveal modular architecture and expand functional coverage in bacteriophage proteins

**DOI:** 10.64898/2026.11.10.765650

**Authors:** Bogna Smug, Krzysztof Szczepaniak, Jade Leconte, Stanisław Dunin-Horkawicz, Rafal J. Mostowy

## Abstract

Bacteriophage proteins evolve through extensive modular exchange, yet the sequence units underlying this process remain poorly defined. Classical domains are plausible evolutionary modules, but they incompletely annotate phage proteins and may not always coincide with the regions conserved across protein families. Here, we constructed 13,035 Evolutionarily Conserved Fragments (ECFs), grouped into 3,167 families, through large-scale HMM–HMM comparisons of proteins from 5,396 RefSeq phage genomes, without imposing predefined domain boundaries. ECFs overlapped substantially with ECOD and Pfam annotations but formed a heterogeneous spectrum comprising subdomains, domain-equivalent units, composite regions and fragments with no counterpart in either resource. Their relationships were frequently nested, indicating that conserved modules occur at multiple organisational scales. ECFs annotated 50% of phage representative proteins — at least doubling the coverage of either database — with the largest gains in structural and infection-related functions, a pattern replicated in independent metagenomic data. ECF families also showed function-dependent patterns of diversity and spatial segregation and provided functional associations for 23,925 protein families lacking detectable Pfam or ECOD domains. In receptor-binding proteins, ECFs subdivided established domains, delineated previously unannotated structural elements and recovered homologous regions where structural similarity was ambiguous. These results show that the evolutionary building blocks of phage proteins do not always coincide with classical domains, highlighting that evolutionary processes act across multiple scales. ECFs provide a complementary, sequence-derived framework for investigating how complex viral proteins are assembled, diversified and functionally organised.

## Introduction

Bacteriophages—viruses that infect bacteria—evolve by mix-and-match assembly rather than clonal descent. Early comparative analyses of complete phage genomes revealed their extraordinary mosaicism, with distantly related viruses sharing conserved modules in patterns inconsistent with simple vertical inheritance ^1,2^. Such mosaicism is driven by diverse recombination mechanisms, including both homologous and non-homologous recombination, which facilitate the exchange of functionally related blocks of genes between different viruses ^3,4,5,6,7^. Over evolutionary time, this pervasive genetic shuffling connects phage lineages into a global gene-sharing network through continuously exchanged modules ^8,9^. Importantly, mosaicism is not confined to genomes but extends to individual proteins, which themselves show evidence of sequence modularity often interpreted as the result of protein-domain exchange ^3,4^. Large-scale analyses of within-protein domain mosaicism have provided support for the fact that protein-sequence modularity can emerge through recombinational exchange of entire domains ^10^, and such exchanges have perhaps been most extensively documented and structurally characterised in receptor-binding proteins ^11^ and endolysins ^12^. The notion that phage protein evolution is driven by domain shuffling is well motivated: domains can fold, function and evolve as semi-autonomous units, making them plausible modules to be gained, lost and exchanged during evolution.

Nevertheless, our understanding of phage protein evolution remains incomplete for two main reasons. The first reason is that detecting domains in phage proteins remains difficult. Resources such as ECOD, CATH and SCOP, which describe the domain universe and define structural, topological, and/or evolutionary relationships between folds, were built largely from proteins of cellular organisms^13,14,15^. Even TED—a recent, massive expansion of the known domain space—remains dominated by cellular proteins, because it inherits the taxonomic composition of AlphaFold DB/UniProt ^16,17^. At the same time, structural modelling of viral proteins has demonstrated that they occupy regions of structural space that are poorly represented in cellular protein resources ^18^. Consequently, many phage proteins lack domain annotations entirely or receive only partial assignments ^19,10,20^. Current domain databases therefore remain insufficient for resolving the full modular architecture of phage proteins and, by extension, for understanding how these proteins evolve.

The second and deeper problem, however, is that the unit of evolution does not always coincide with the domain. Evolutionary units are typically identified through genealogical homology. Sequence similarity provides the most direct evidence of homology because non-trivial similarity across distant proteins is far more likely to reflect shared ancestry than independent emergence ^21^. When conserved units are detected this way – *de novo*, from sequence, without imposed boundaries – they form a spectrum rather than clean domains: highly conserved sub-regions surface as subdomain fragments; co-evolving domains surface as composite units known as superdomains ^22,23,24^. Independent evidence points in the same direction. Large-scale sequence comparisons of known protein domains reveal that sub-domain segments are widespread, shared between proteins of completely independent evolutionary origin, and likely predate the domains that contain them^21,25,26,27^. Evolutionary building blocks may therefore be nested and overlapping, rather than discrete, mutually exclusive units the domain framework assumes ^28^. Whether phage protein building blocks fall above or below the domain level has not been tested. Answering this question, and compensating for the limited domain coverage of phage proteins, requires an empirical catalogue of conserved units compared against established domain resources as a reference frame.

To address this challenge, we built a resource called Evolutionarily Conserved Fragments (ECFs): 13,035 empirically derived, conserved sequence units, identified through large-scale HMM-HMM comparisons across 5,396 reference phage genomes, without imposing pre-existing boundaries. To characterise whether ECFs correspond to known domains, fall above or below them, or represent sequence space with no curated counterpart, we compared them against two complementary resources — ECOD, the primary structural reference for domain evolution, and Pfam, which defines domain families from sequence rather than structure. We then assessed both the extent of overlap and the nature of the discrepancies. Finally, we evaluated the functional coverage of ECFs, Pfam and ECOD databases across RefSeq bacteriophages and within an independent metagenomic dataset. The conserved units of phage protein modularity formed a heterogeneous spectrum — whole domains, subdomains, composite regions, and genuinely novel fragments — that current structural and sequence-based resources captured only in part. We present ECFs as a phage-specific, sequence-first resource, complementary to existing domain databases. Above all, ECFs provide a framework for dissecting modularity below the domain level in bacterial viruses and beyond, and for studying how viral proteins evolve and acquire new functions through modular exchange.

## Results

### ECFs define recurrent units of phage protein architecture

To identify Evolutionarily Conserved Fragments (ECFs) in phage proteins, we developed a large-scale, profile-based fragment detection pipeline inspired by earlier de novo domain detection approaches ^22^ but exploiting the sensitivity of HMM-HMM comparison ^29^. The approach is described in detail in Methods and summarised in Figure 1A. Briefly, we predicted protein sequences from 5,396 RefSeq phage genomes, clustered them into 148,010 representative groups, and built a profile HMM (rHMM) for each group. An all-vs-all HMM comparison of rHMMs was then used to identify recurrent alignment regions within each query protein; these were clustered by position to define query-level fragment candidates (qECFs), which were filtered based on overlap consistency and coverage statistics calibrated against high-confidence Pfam assignments. The resulting 121,594 qECFs were converted into HMM profiles and compared exhaustively against one another; pairs with strong reciprocal coverage were grouped by graph-based clustering into 13,035 Evolutionarily Conserved Fragments (ECFs). ECF profiles were then searched against structural (ECOD) and sequence-based (Pfam) domain databases, and ECFs were further grouped by similarity into 3,167 higher-order ECF families (EFs).

**Figure 1.**
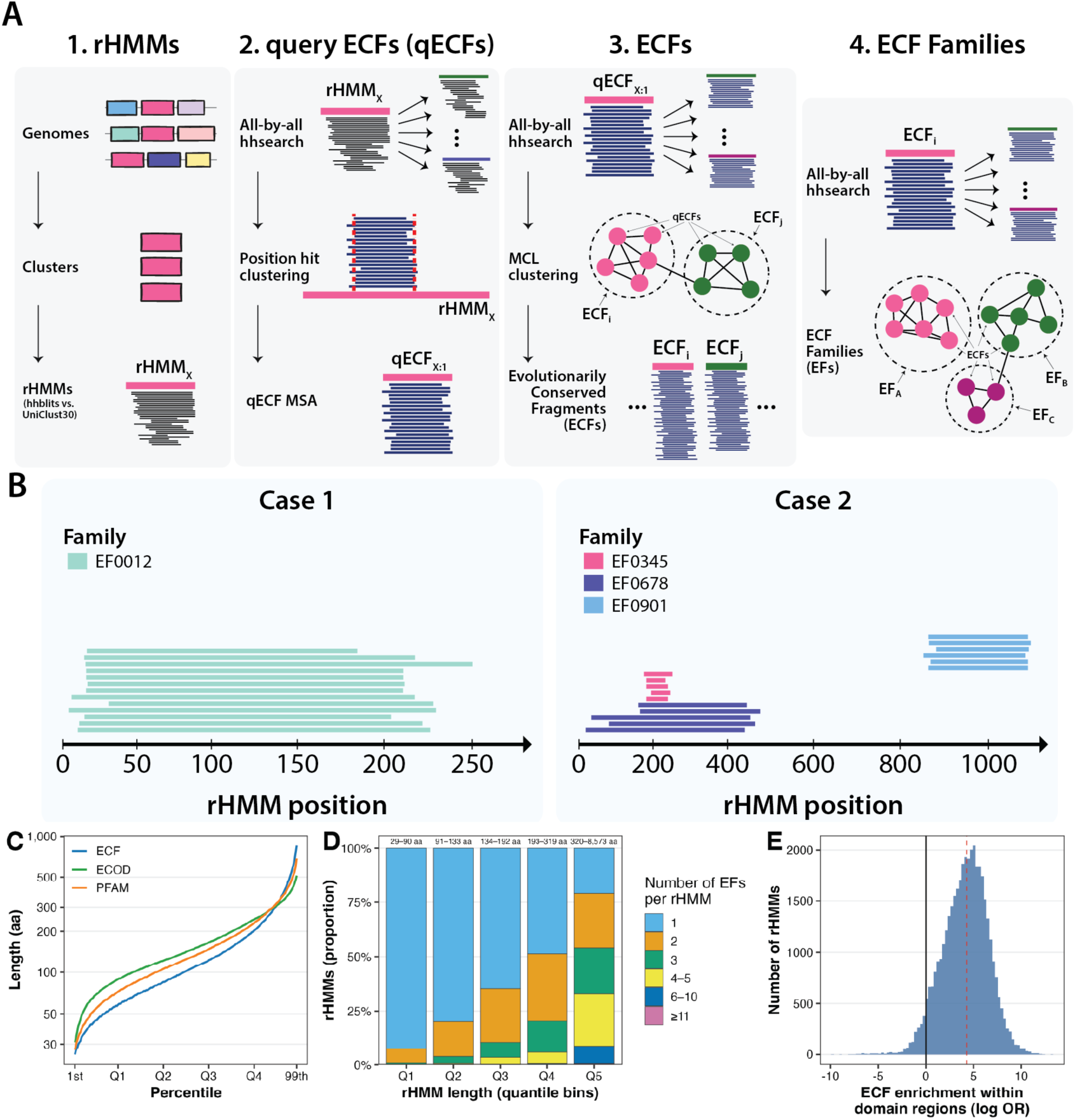
Construction and basic properties of Evolutionarily Conserved Fragments. (A) Overview of the ECF detection pipeline. Protein sequences predicted from 5,396 RefSeq bacteriophage genomes were clustered into representative groups, each represented by a profile HMM (rHMM). An all-by-all HMM comparison of rHMMs was used to identify recurrent alignment regions within each query protein; these were clustered by position to define query-level fragment candidates (qECFs), each summarised as a multiple sequence alignment. qECF profiles were then compared exhaustively against one another, and pairs with strong reciprocal coverage were grouped by MCL clustering into Evolutionarily Conserved Fragments (ECFs). Finally, overlapping ECFs are assigned into Families (EFs). (B) Synthetic examples of ECF hits mapped onto query rHMMs. When query profiles (rHMMs) are searched against all 13,035 ECF profiles, significant hits are projected onto the query and coloured by ECF family (EF). Case 1 shows one dominant EF spanning most of the protein; Case 2 shows three EFs arranged in separate and partially nested regions. (C) Length distributions of ECF (blue), ECOD (green) and Pfam (orange) entries of at least 20 amino acids. Lengths are plotted on a log scale against percentile rank. (D) Number of distinct ECF families detected per rHMM across protein length quintiles (Q1–Q5; length ranges indicated above each bar). (E) Enrichment of ECF coverage within domain-annotated regions. For each rHMM containing both ECF and domain (Pfam or ECOD) hits, we calculated the log odds ratio of ECF coverage inside versus outside annotated domains, applying a Haldane–Anscombe correction of 0.5 to each contingency-table cell. The solid line marks independence (log OR = 0); the dashed line marks the median (log OR = 4.27). Positive values indicate enrichment within domain-annotated regions.

Figure 1B provides a schematic illustration of how ECFs can be visualised within a query protein. The examples are synthetic but reflect patterns commonly observed in the data. First, a query sequence alignment is converted into an HMM profile and searched against the 13,035 ECF profiles following the HHpred framework ^30^. Then, significant matches are projected onto the query sequence and coloured according to their EF assignment. In the first example, most of the protein is covered by ECFs from a single EF; in the second, hits from three EFs form a more complex nested and composite arrangement. Although these maps resemble searches against domain databases like ECOD or CATH, ECFs are defined differently: they are domain-agnostic units inferred solely from recurrent sequence conservation across phage protein families.

We next examined ECF size and reuse. Individual ECFs were detected in a median of 23 rHMMs, with a long tail extending to several hundred occurrences, indicating that most are recurrent but not ubiquitous (Supplementary Figure S1). While individual ECFs recurred, the catalogue as a whole was diverse: most EFs were small, with 1,449 EFs consisting of a single ECF and further 1,211 containing between two and five ECFs (Supplementary Figure S2). ECFs had a median length of 102 amino acids, compared with 142 for ECOD domains, 124 for Pfam domains, and 159 for full rHMMs (Supplementary Figure S3). They were therefore typically shorter than established domains and their parent protein families, but remained broadly domain-sized. Their length distribution was also wider than those of ECOD and Pfam, with an excess of both short and long fragments (Figure 1C), consistent with ECFs capturing subdomain units as well as composite regions extending beyond individual domain boundaries. Longer rHMMs contained a greater diversity of ECF families. After dividing rHMMs into length quintiles, the shortest proteins (29–90 amino acids) were dominated by a single EF, whereas the longest (321–8,573 amino acids) contained a median of three and frequently six or more EFs (Figure 1D). Nine rHMMs contained at least 20 distinct EFs and had a median length of 4,422 amino acids, highlighting exceptionally complex architectures. Thus, EF diversity increases with protein length, consistent with their resolving genuine architectural complexity rather than partitioning proteins into a fixed number of fragments.

Finally, we asked whether ECFs behaved as structured conserved units rather than arbitrary sequence fragments. We first compared the positions of ECF hits along rHMMs with those of Pfam and ECOD hits, plotting the relative start and end of each hit within its protein (Supplementary Figure S4). Although ECFs produced a denser pattern, consistent with their greater number and shorter average length, their positional distribution was qualitatively similar to that of established domains. We therefore tested whether ECFs were preferentially located within domain-annotated regions. For each rHMM containing at least one ECF hit and at least one Pfam or ECOD assignment, we reduced all hit coordinates to non-overlapping intervals and classified residues as covered by both an ECF and a domain, by an ECF only, by a domain only, or by neither. We then quantified the association between ECF and domain coverage using a log odds ratio, applying a Haldane–Anscombe correction of 0.5 to each category (so the log odds ratio stays defined even when a count is zero). ECFs were strongly enriched within domain-annotated regions, with a median log odds ratio of 4.27 (Figure 1E). ECFs were therefore not randomly distributed fragments: they shared important positional properties with Pfam and ECOD domains and were strongly associated with them, although the precise nature of this relationship remains unresolved and is examined in the next section.

### ECFs reveal non-obvious links between distant parts of the phage protein space

Having established that ECFs behave as structured conserved units rather than arbitrary sequence fragments, we next asked how they relate to each other using all high-confidence ECF–ECF (HMM-HMM) alignments. For each ECF, we summarised its alignments by the median query and subject coverage and plotted these values in coverage space (Figure 2A). The resulting distribution was strongly L-shaped, with ECFs concentrated away from the diagonal and towards combinations of high coverage in one sequence and low coverage in the other. The same pattern was evident amongst individual alignments and when connections were summarised by their mean coverage, indicating that it was a general property of the ECF catalogue rather than an artefact of a small number of highly connected ECFs (Supplementary Figure S5). Such asymmetry is expected when one conserved unit is nested within another: high query and low subject coverage identifies a shorter, **inset** ECF contained within longer ones, whereas the converse identifies **container** ECF (i.e., one containing other ECFs). Consistent with this interpretation, coverage asymmetry was related to ECF length, with shorter ECFs tending to behave as insets and longer ECFs as containers; insets (qcov > scov) were more common than containers (qcov < scov; see Supplementary Figure S6). The ECF–ECF network therefore exhibited a hierarchical containment structure, in which shorter inset ECFs were frequently embedded within larger, conserved units (containers).

**Figure 2.**
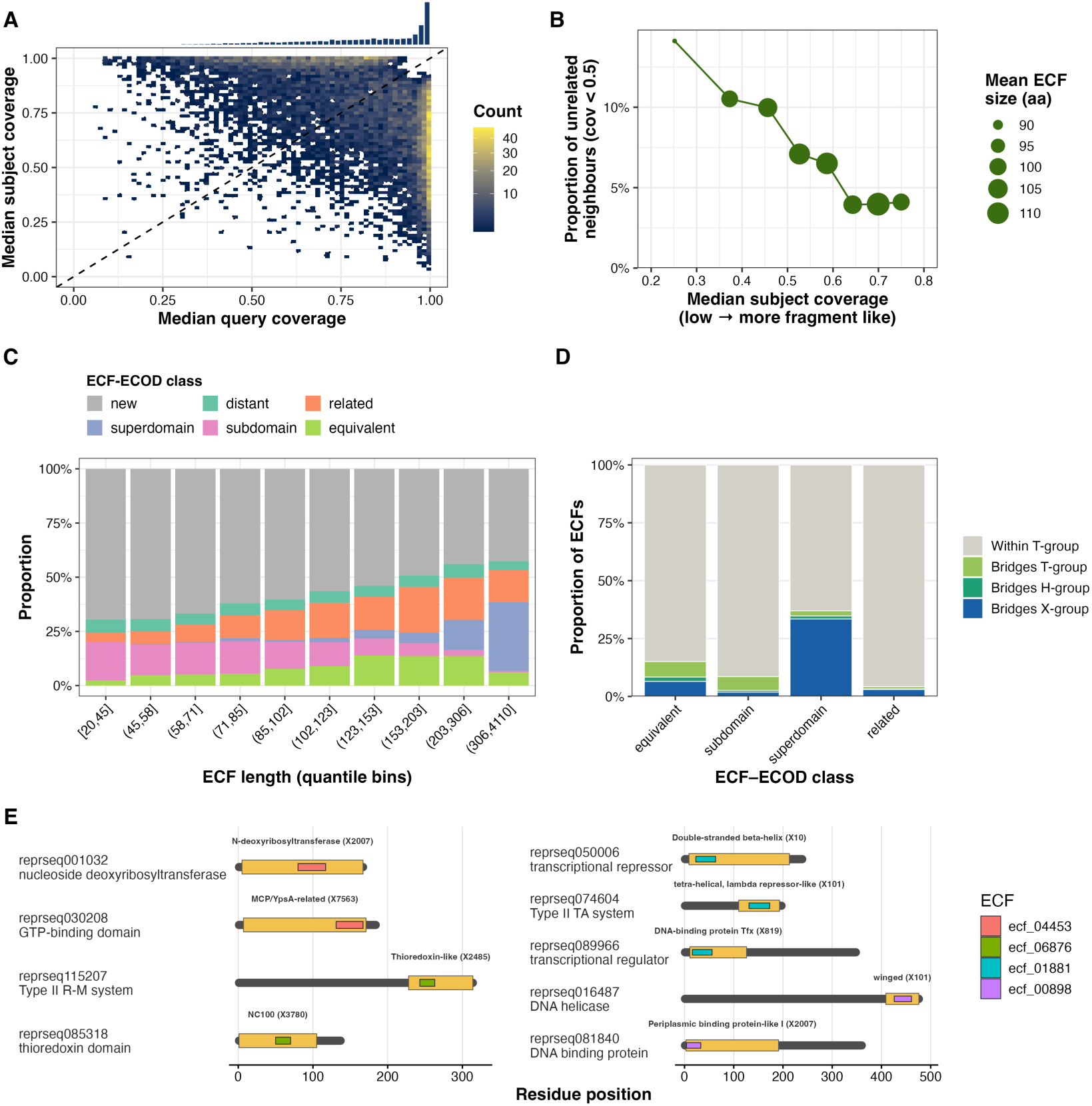
Hierarchical and domain-relationship structure of ECFs. (A) Per-ECF median query and subject coverage across all high-confidence ECF–ECF connections (probability ≥ 99%), shown as a 2D density heatmap with a marginal histogram of median query coverage. The dashed line indicates qcov = scov. (B) Proportion of unrelated container pairs (maximum pairwise coverage < 0.5), plotted against the degree of containment of their shared inset ECF, quantified as its maximum subject coverage across the two inset–container alignments. Values were grouped into equal-size deciles (left to right: stronger to weaker containment). Point size indicates the mean ECF length within each decile. (C) Proportional distribution of ECF–ECOD domain assignment classes across ten equal-frequency ECF length bins. Each ECF was assigned the highest-priority class meeting the following criteria: equivalent (probability ≥ 95%, e-value ≤ 0.01, qcov ≥ 0.8, scov ≥ 0.8); subdomain (probability ≥ 95%, e-value ≤ 0.01, qcov ≥ 0.9, scov ≤ 0.6); superdomain (probability ≥ 95%, e-value ≤ 0.001, qcov ≤ 0.6, scov ≥ 0.9); related (probability ≥ 95%, not meeting the above); distant (no hit meeting these thresholds); new (no confident hit to ECOD at any threshold). (D) Proportion of ECFs in each ECOD assignment class (equivalent, subdomain, superdomain, related) falling into four ECOD-bridging categories — within a single T-group, bridging T-groups, bridging H-groups, or bridging X-groups — based on the number of distinct ECOD groups matched with at least two confident hits per group. Categories were assigned from distinct ECOD groups represented by at least two confident hits. (E) Representative example proteins for four hand-selected ECFs (ecf_00043, ecf_00108, ecf_01881, ecf_04622), each shown on one representative protein per ECOD X-group it matches. Grey bars represent protein length; gold rectangles indicate ECOD domain boundaries; coloured rectangles indicate ECF boundaries (one colour per ECF, shared across panels). Domain name and X-group identifier are annotated above each ECOD hit.

We next asked whether inset ECFs simply represented the most conserved subregions of otherwise related proteins, or whether they acted as reusable modules shared between unrelated sequence contexts. For each inset ECF, we therefore examined pairs of ECFs that contained it and measured how well those containers aligned to one another at different degrees of containment, quantified as the maximum subject coverage of the inset ECF against the two containers. If the inset ECF belonged to a single conserved parent unit, its containers should themselves be closely related; by contrast, an inset reused across different proteins could connect containers with little or no detectable similarity outside the shared region. The proportion of unrelated container pairs increased with the degree of containment, reaching approximately 15% among the most strongly contained ECFs (Figure 2B). Near-zero or zero similarity between containers was also observed across the full range of containment values (Supplementary Figure S7). At least some ECFs therefore did not merely mark conserved cores within a single protein family, but bridged otherwise unrelated conserved units, consistent with their reuse as modular evolutionary building blocks.

We next compared ECFs directly with established protein domains, assigning each to one of sex classes according to the geometry of its best ECOD match: ‘equivalent’, ‘subdomain’, ‘superdomain’, ‘related’, ‘distant’ or ‘new’ (see Methods). These classes showed a strong dependence on ECF length (Figure 2C), with shorter ECFs more often classified as subdomains and longer ECFs increasingly classified as superdomains or related regions. The same qualitative pattern was recovered against Pfam (Supplementary Figure S8), and the two databases showed substantial agreement in their assignments (Supplementary Figure S9). Notably, however, the largest class comprised ECFs with no detectable counterpart in either resource. This fraction decreased as the search threshold was relaxed, but remained 42% at an e-value of ≤ 0.1 and 15% even at the highly permissive threshold of ≤ 10 (Supplementary Figure S10). Proteins carrying at least one ‘new’ ECF were significantly enriched for phage-specific structural functions (head-and-packaging, connector, tail) but also for morons, lysis and kinase functions (Supplementary Figure S11). Enriched Pfam families showed a similar bias, with many corresponding to terminase, portal, capsid and tail proteins, indicating that unmatched ECFs concentrate in coherent, annotation-poor structural contexts rather than representing spurious fragments (Supplementary Figure S11). Thus, ECFs span a heterogeneous spectrum of relationships to known domains, from subdomain fragments to whole domains and composite regions, while also recovering conserved, functionally coherent units that remain outside current domain annotations.

Given the hierarchical structure of the ECF network, we next hypothesised that ECFs might reveal non-obvious links between established domain architectures. To test this, we used the hierarchical classification of ECOD, in which domains are grouped by topology (T-groups), sequence-level homology (H-groups), and broader structural homology (X-groups). For each ECF with multiple confident ECOD matches, we determined whether its hits remained within a single T-group (‘Within T-group’) or extended across distinct T-groups (‘Bridges T-groups’), H-groups (‘Bridges H-groups’), or X-groups (‘Bridges X-groups’). Figure 2D shows the distribution of these categories across ECF–ECOD classes. Most ECFs in every class were restricted to a single T-group, as expected for fragments closely matching an established domain. However, all classes also contained ECFs that linked domains across higher levels of the ECOD hierarchy, including distinct H-groups, for which homology is supported only structurally, and distinct X-groups, for which ECOD reports no evidence of homology from structural comparison. Bridging across X-groups was most frequent among ECFs classified as superdomains, consistent with their greater likelihood of spanning multiple domains. Importantly, the presence of H- and X-group bridging among equivalent, subdomain and related ECFs indicates that conserved sequence fragments can recur within domains assigned to evolutionarily distant lineages, in some cases without detectable homology at either the sequence or structural level. Thus, the mismatch between ECF and domain boundaries is not purely geometric: some ECFs reveal connections between otherwise distinct domain architectures.

To illustrate these relationships directly, we selected four ECFs that matched domains from at least two ECOD X-groups and occurred in proteins with distinct functional annotations (Figure 2E). For each ECF, representative proteins from different X-groups are shown with the positions of the ECF and ECOD assignments mapped onto their sequences. In each case, the same conserved fragment occurred within proteins carrying different ECOD domains and broader architectures. These examples provided concrete instances of the relationships inferred across the catalogue, showing how an ECF can represent a conserved module shared between proteins that are not grouped together by conventional domain classifications.

### ECFs expand functional annotation coverage of phage proteins

The large fraction of ‘new’ ECFs, together with their enrichment in phage-specific functions, suggested that ECFs might recover substantially more of the phage protein space than existing domain resources. We hypothesised that, because ECFs were derived empirically from viral sequences rather than from cellular protein databases, they would provide broader annotation coverage, particularly for structural and phage-specific functions. To test this, we searched all 148,010 rHMMs against ECF, ECOD, and Pfam under identical detection parameters (probability ≥ 95%, e-value ≤ 0.1, minimum hit length ≥ 20 aa, subject coverage ≥ 70%) and classified each rHMM as annotated if it received at least one hit. The largest single category comprised rHMMs annotated by none of the three databases (71,488; Figure 3A), underscoring the large unannotated fraction of phage protein space. ECF-only annotations formed the second largest category (34,549), well ahead of proteins annotated exclusively by Pfam (1,103) or ECOD (410). Across PHROG functional categories, ECF coverage exceeded both Pfam and ECOD in nearly all classes (Figure 3B), with the largest gap for tail proteins, where ECF coverage exceeded 95% against 56% for Pfam and 42% for ECOD (see also Supplementary Figure S12). The databases performed most comparably for metabolic functions — including DNA/RNA metabolism and integration and excision — though ECF coverage remained higher or equivalent throughout. Even for proteins of unknown function, ECFs (34%) substantially outperformed Pfam (9%) and ECOD (7%). Overall, ECFs annotated 50% of rHMMs, at least twice the coverage of Pfam (25%) or ECOD (20%).

**Figure 3.**
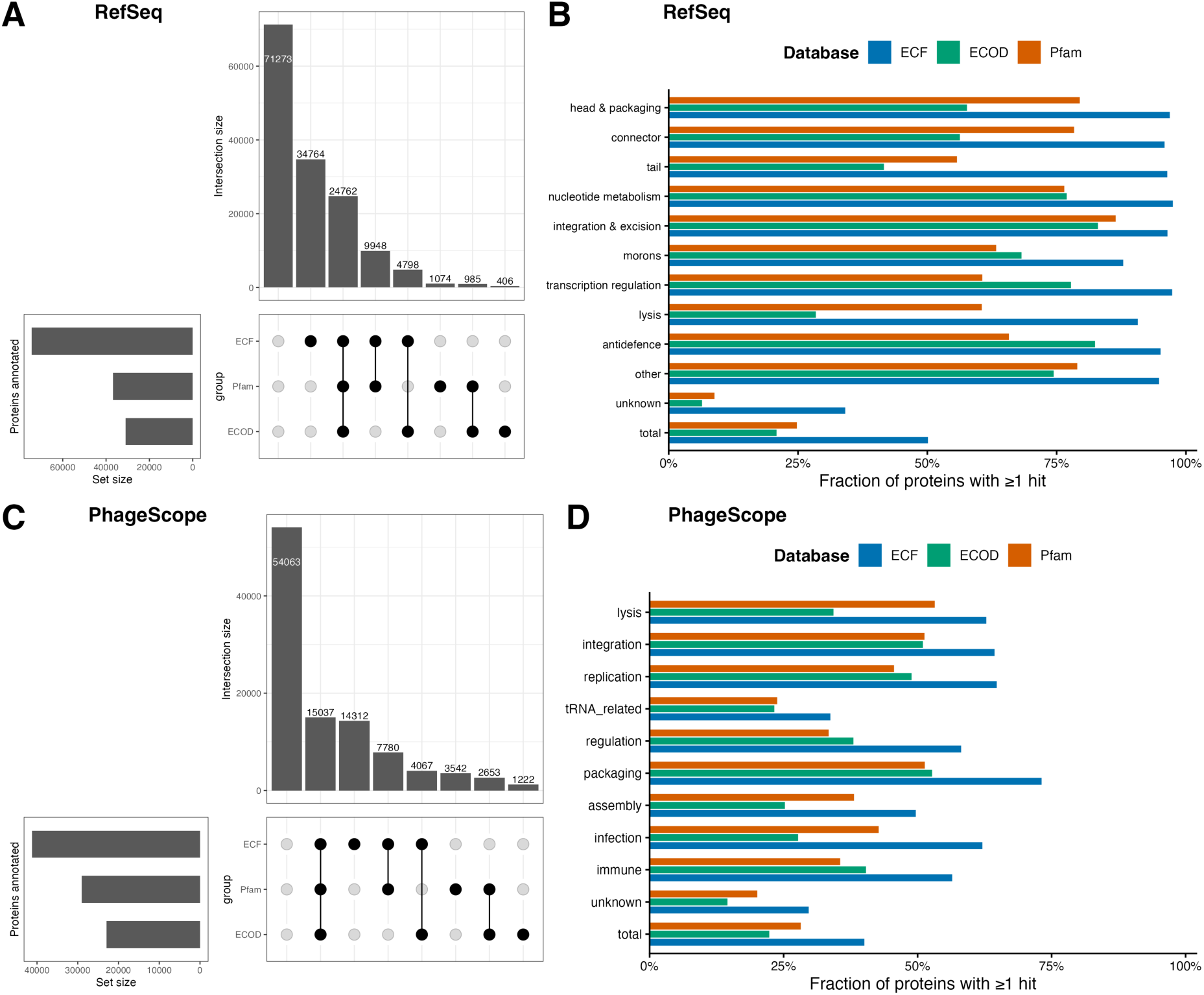
Annotation coverage of phage proteins by ECF, ECOD, and Pfam. (A) UpSet plot comparing proteins in the RefSeq representative sequence dataset annotated by ECF, Pfam and ECOD. Horizontal bars (left) show the total number of rHMMs annotated by each database; vertical bars show the size of each exclusive intersection. (B) Fraction of rHMMs with at least one hit from each database, broken down by PHROG functional class, lack thereof (‘unknown’) or across the full dataset (‘total’). (C) UpSet plot as in (A) for 102,676 proteins from 2,000 PhageScope phage genomes not represented in the RefSeq training set. (D) Fraction of the PhageScope proteins with at least one hit from each database, broken down by PhageScope functional category. In all panels, ECF, ECOD, and Pfam hits were detected using identical thresholds (probability ≥ 95%, e-value ≤ 0.1, minimum hit length ≥ 20 aa, domain coverage ≥ 70%).

One concern was that the ECF advantage could reflect circularity, as ECFs were built from the same rHMMs against which coverage was measured. We therefore repeated the analysis in an independent set of 2,000 phage genomes sampled from PhageScope, pre-filtered to exclude any genome present in GenBank or RefSeq, and searched their 102,676 proteins against ECF, ECOD and Pfam under identical thresholds. The unannotated fraction was again the largest single category (54,174; Figure 3C), and proportionally larger than in RefSeq, consistent with the greater sequence novelty of metagenomic phage genomes. Proteins annotated by all three databases ranked second (14,960), followed closely by ECF-only annotations (14,201); Pfam-only (3,492) and ECOD-only (1,263) proteins again formed small minorities. Across functional categories, ECF coverage exceeded both ECOD and Pfam in all classes, with the largest gaps for infection-related (ECF 62% vs. ECOD 28% vs. Pfam 43%) and assembly proteins (ECF 50% vs. ECOD 25% vs. Pfam 38%; Figure 3D). The databases converged most closely for integration and replication, where Pfam (51%) and ECOD (51%) approached ECF coverage (64%), consistent with these categories harbouring enzymatic proteins with well-characterised cellular homologues. Overall, ECFs provided substantially broader annotation coverage than either curated database, with the greatest gains in structural and phage-specific functional classes.

### ECF families organise phage proteins into function-dependent architectural regimes

Having shown that ECFs extend beyond established domain definitions and recover otherwise hidden regions of phage protein space, we next asked whether their arrangement within proteins could reveal non-obvious patterns behind the evolutionary protein architecture across different functions. In particular, we examined whether the number and spatial organisation of ECF families capture distinct patterns of modular evolution. The schematic examples in Figure 1B already suggest that ECFs detected within a single query could distribute into unique ECF Families (EFs), and that the EFs could vary in their degree to occupy separate or overlapping sequence regions. The first measure reflects EF diversity while the second one reflects EF segregation.

EF diversity, defined as the number of distinct EFs detected in a protein (`n.fams`), increased with the total number of ECF hits, as expected (Figure 4A). However, the relationship was broad rather than deterministic: proteins with similar numbers of ECF hits often differed substantially in the number of EFs represented. Exceptionally long proteins contributed to the upper tail of the distribution, but the same loose relationship remained after restricting the analysis to proteins shorter than 1,000 amino acids (Supplementary Figure S13). Thus, EF diversity was not simply a direct consequence of detection depth.

**Figure 4.**
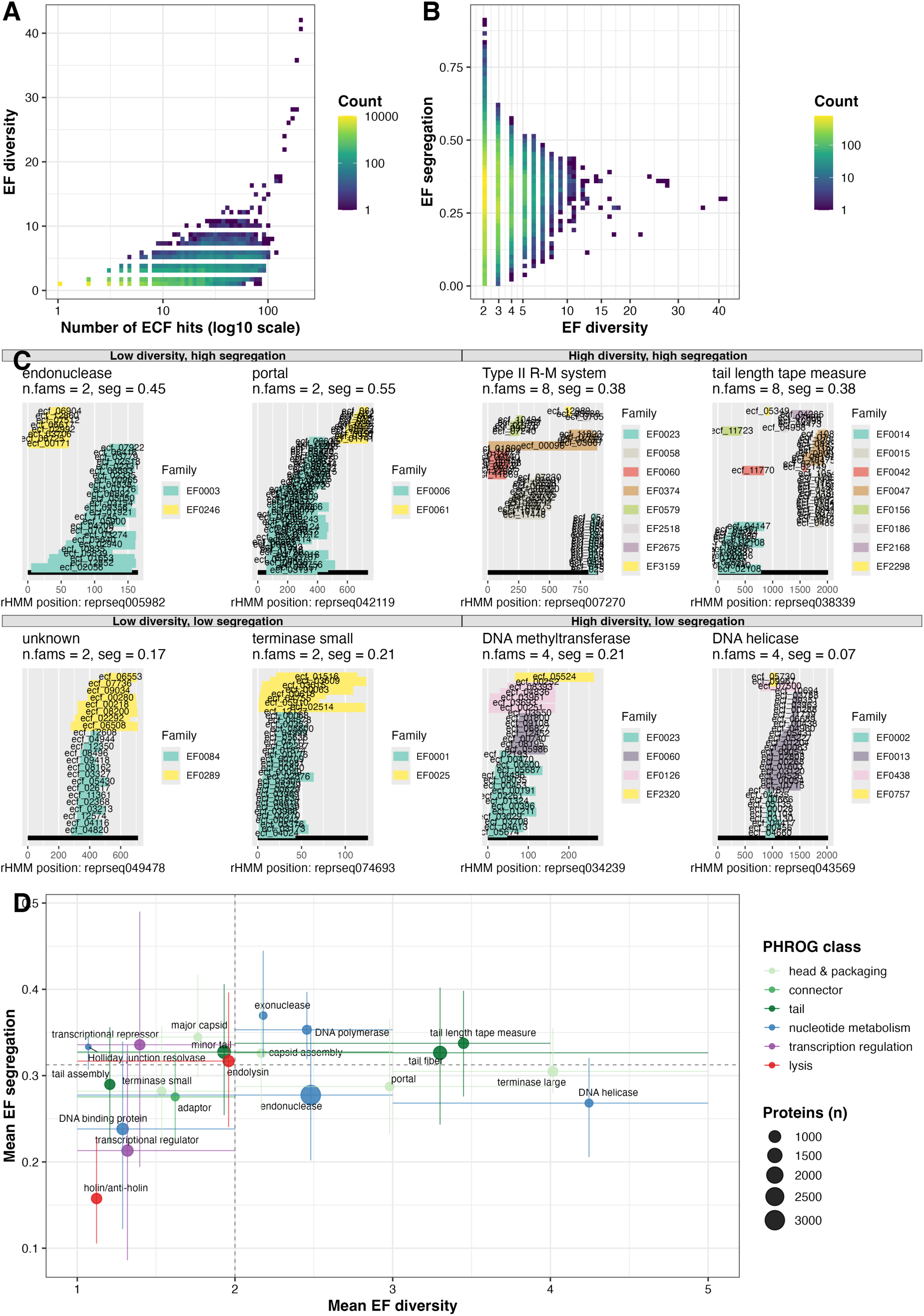
ECF family architecture of phage proteins. (A) EF diversity against detection depth. Binned density of the number of distinct EFs (`n.fams`) against the total number of ECF hits (`n.ecf.hits`, log_10_ scale) across all rHMMs with non-zero ECF coverage. Bin counts are shown on a log_10_ viridis colour scale; the dashed line marks *y* = *x*, the upper bound attainable if every hit came from a different family. (B) EF diversity against EF segregation. Binned density of the number of EFs (square-root scale) against the barycentric segregation index, across rHMMs carrying at least two EFs. Colour scale as in (A). (C) Eight representative rHMMs spanning the four diversity–segregation combinations, arranged to mirror (B): the upper block shows high-segregation proteins and the lower block low-segregation proteins, with low-diversity examples on the left of each block and high-diversity examples on the right. Each subpanel plots individual ECF hit intervals against position in the query rHMM, coloured by EF, and is titled with the PHROG function of the protein together with its `n.fams` and segregation values. (D) Functional categories in the EF diversity–EF segregation plane. The twenty PHROG functions with the most annotated rHMMs are plotted by their mean number of EFs (x-axis) and mean barycentric segregation index (y-axis); only functions with at least 30 proteins contributing to each axis were retained. Point size indicates the number of proteins carrying the function and colour indicates PHROG class. Horizontal and vertical bars span the interquartile range of each measure. Dashed grey lines mark the median across all qualifying functions.

We next quantified EF segregation as the spatial separation of family barycentres along the protein sequence (`segregation.bary`). Diversity and segregation were largely independent: proteins with the same number of families spanned a broad range of segregation values, particularly at low and intermediate diversity (Figure 4B). The attainable segregation range narrowed at high diversity, as expected from the geometric constraint of accommodating many distinct families within a finite sequence. Consequently, proteins with identical EF diversity could nevertheless display markedly different internal architectures.

Figure 4C shows representative examples of proteins with various functions with different combinations of EF diversity/EF segregation (high/high, high/low, low/high, low/low). At low diversity, EF families either formed distinct, non-overlapping blocks, as in representative endonuclease and portal proteins, or occupied the same sequence region, as in a small terminase and a protein of unknown function. At high diversity, families formed either spatially separated clusters, as in a Type II restriction–modification system protein and a tail length tape measure protein, or densely interleaved mosaics, as in representative DNA methyltransferase and DNA helicase proteins. These examples show that similar family counts can correspond to fundamentally different modular organisations.

Because segregation assumes that each EF family occupies a reasonably consistent position within a protein, we evaluated within-family positional agreement using average hit overlap (AHO). AHO was high overall, with a median of approximately 0.6, but varied across functional categories (Supplementary Figure S14). The lowest values were concentrated in connector- and tail-associated proteins, including tail length tape measure proteins, where repeated structural units recur at multiple positions. Low AHO in these proteins therefore reflected genuine repeat architecture rather than unstable ECF placement, consistent with the repeated and multi-cluster patterns visible among the representative examples.

We also assessed whether EF diversity and EF segregation adequately summarised a broader set of candidate architecture metrics. To this end, we proposed two additional measures of EF diversity (mean effective number of EFs and number of dominant EF segments) and another measure of EF segregation (overlap-based index; see Methods). All three EF diversity measures were strongly intercorrelated, while the two EF segregation measures were partly correlated (Supplementary Figure S15). Principal component analysis of five rank-transformed metrics identified two dominant components explaining 76% of the total variance (Supplementary Figure S16). EF diversity (`n.fams`) tracked the first component, whereas EF segregation (`segregation.bary`) tracked the second, supporting their use as representative measures of two largely independent architectural dimensions (Supplementary Figure S16).

Finally, we asked whether phage protein functions occupied distinct regions of the EF diversity–EF segregation landscape. The twenty most protein-rich functional categories showed systematic but continuous differences in this 2D space (Figure 4D). For example, holin/antiholin and DNA-binding proteins were concentrated towards low diversity and low segregation, whereas capsid proteins and transcriptional repressors combined low diversity with greater segregation. Several functions showed consistently high diversity but differed in how their EF families were arranged: exonucleases, tail fibres and tail length tape measure proteins tended towards high segregation, while endonucleases, helicases, and large terminases occupied regions of lower segregation. Although individual proteins varied substantially within each function and neighbouring functional distributions overlapped, their average positions remained distinct. EF diversity and segregation therefore capture function-associated differences in protein architecture while also revealing considerable within-function heterogeneity, consistent with multiple modular solutions to related biological roles.

Together, these analyses show that ECF families are organised within phage proteins along two largely independent dimensions: how many distinct families are present and how strongly those families are spatially separated. These dimensions capture architectures ranging from clean two-block arrangements to highly interleaved multi-family mosaics and vary systematically with protein function, indicating that ECF distributions encode architectural information beyond conventional domain annotation. This information may in turn reflect differences in how protein functions were assembled and diversified: spatially segregated families are compatible with the combination of blocks of different evolutionary origin, whereas overlapping families are compatible with diversification within a shared block. However, structural complexity, internal repeats and protein length can generate similar patterns, so disentangling these processes will require explicit evolutionary analyses.

### ECFs provide functional rescue of domain-dark proteins

Given the complementary nature of ECFs with respect to domain databases, the elevated coverage and non-trivial insight into the architectural design of diverse functions, we hypothesised that ECFs could provide functional information for proteins that ECOD and Pfam fail to annotate entirely. We termed such proteins *domain-dark*: rHMMs carrying at least one confident ECF hit (probability ≥ 95%, ECF coverage ≥ 70%, hit length ≥ 20 aa) but lacking any Pfam or ECOD hit at lenient detection thresholds (e-value ≤ 0.1, hit length ≥ 20 aa). This conservative definition yielded 23,925 domain-dark proteins. The question was whether, among these proteins, carrying a specific ECF family (EF) made a protein significantly more likely to belong to a particular functional class. Such an association would suggest that the EF can serve as a functional signature for domain-dark proteins.

To test this, we restricted the analysis to the 2,959 domain-dark proteins (12%) with a confident PHROG annotation (best hit, score ≥ 0.5), because enrichment can only be measured against known functions. For each EF detected in these proteins and each of the 25 most frequent functions among them, we constructed a 2 × 2 contingency table (family present or absent × function present or absent), using all annotated domain-dark proteins as the background, and applied a one-sided Fisher’s exact test. Pairs with fewer than five domain-dark proteins carrying both the family and the function were not tested. *P*-values were corrected across all tested pairs using the Benjamini–Hochberg procedure, and enrichments were retained at FDR ≤ 0.01. This tests whether EFs are informative about function, which is the prerequisite for assigning function to the remaining unannotated domain-dark proteins.

The number of significantly enriched EFs varied widely across the 25 tested functions, from one (spanin, anti-CBASS) to 54 (tail fibre), and increased with the diversity of EFs present in domain-dark proteins of a given function (Figure 5A). Tail fibre had both the highest EF diversity (number of unique EFs) and the most enriched EFs, followed by minor tail (30), tail length tape measure (26), membrane protein (16) and endolysin (14). Every tested function had at least one enriched EF, with the fewest found for spanin, anti-CBASS, DNA binding protein and anti-repressor (one or two each). Because both quantities increase with the number of domain-dark proteins carrying a given function, this trend partly reflects statistical power. Nevertheless, the enrichments were dominated by tail-associated functions, which are also those for which ECF coverage most exceeded that of Pfam and ECOD (Figure 3B). Functional signal from EFs was therefore most abundant in the structurally diverse, phage-specific protein classes that existing domain databases covered least well.

**Figure 5.**
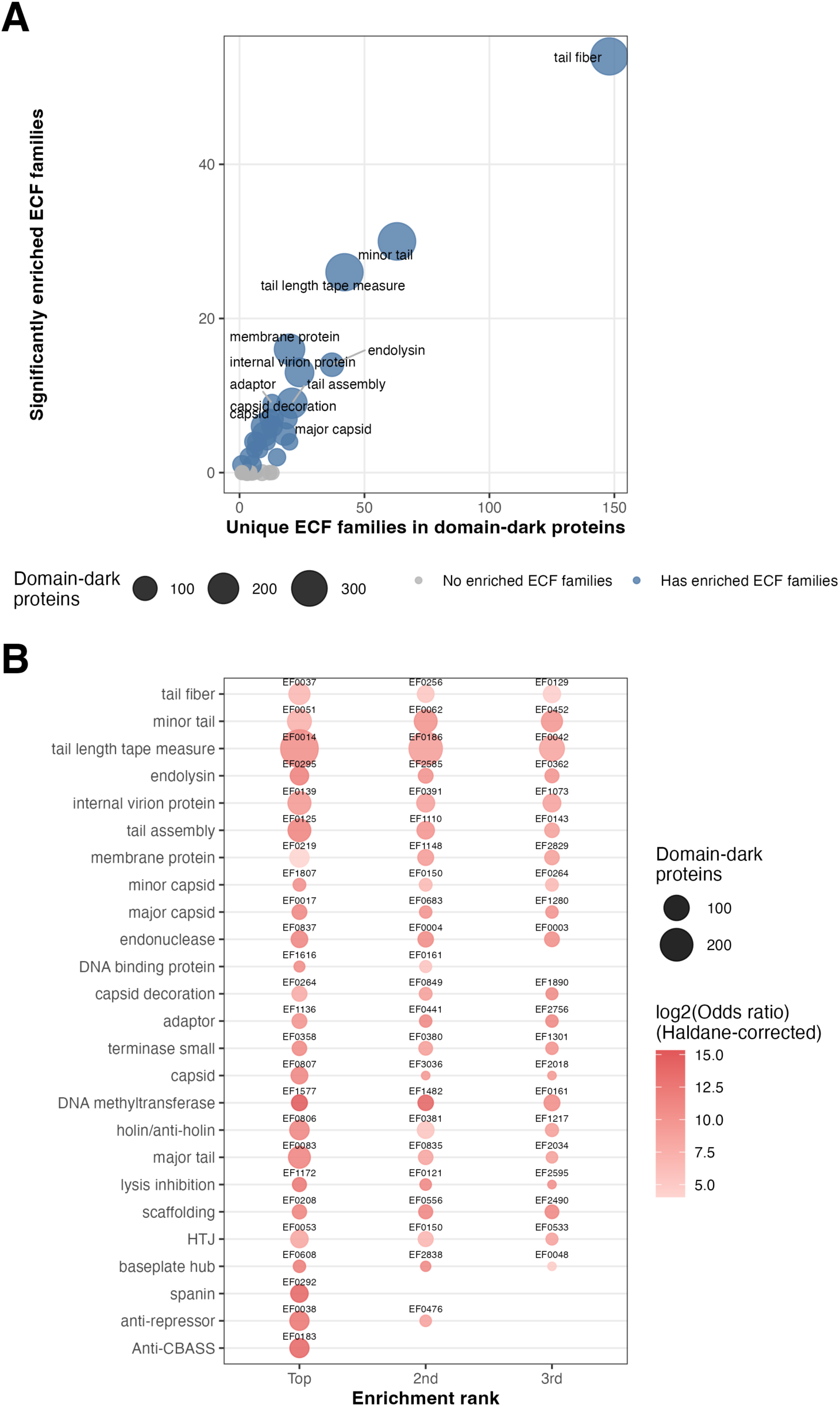
ECFs provide functional rescue of domain-dark proteins. (A) For each of the 25 most frequent PHROG functions among annotated domain-dark proteins (*n* = 2,959), the number of unique ECF families (EFs) detected in domain-dark proteins of that function (x-axis) is plotted against the number of significantly enriched EFs (y-axis; one-sided Fisher’s exact test, Benjamini–Hochberg FDR ≤ 0.01). Point size indicates the number of annotated domain-dark proteins with that function. Domain-dark proteins were defined as rHMMs with at least one confident ECF hit but no Pfam or ECOD hit at lenient detection thresholds (see text). (B) Up to three significantly enriched EFs per function, ranked by *P*-value (ties broken by the total number of domain-dark proteins carrying the EF) and shown from top to bottom in order of decreasing EF diversity in panel A; functions with fewer than three significant EFs show fewer points. Point size indicates the number of domain-dark proteins carrying both the EF and the function; colour indicates the log_2_ odds ratio, with a Haldane–Anscombe correction of 0.5 added to each cell of the contingency table so that the ratio remains finite when a cell is zero (display only; significance is based on the uncorrected test). EF labels are shown above each point.

Finally, we examined the enriched EFs individually (Figure 5B). If an EF is to serve as a functional signature, it should occur in proteins of one functional class and not in others. For each function, Figure 5B therefore shows the three EFs with the most significant enrichment (lowest p-value, with ties broken by the number of domain-dark proteins carrying both the family and the function). The associations were largely exclusive: in 148 of the 242 significant EF–function pairs, every annotated domain-dark protein carrying the EF belonged to the same function. Exclusivity was not restricted to small families, as the top-ranked EFs combined it with many member proteins. For example, EF0125 occurred in 74 domain-dark proteins annotated as tail assembly, none with a different function, and a further 11 domain-dark proteins carrying it lacked a confident functional call and could therefore inherit this assignment. Together, these results indicate that in domain-dark proteins, where Pfam and ECOD provide no information, membership of a specific EF is often a strong and frequently exclusive indicator of functional class, supporting the use of EFs as functional signatures.

### ECFs reveal modular architecture in receptor-binding proteins beyond domain annotations

Previous sections established that ECFs recover homologous sequence modules, expand functional annotation and reveal function-dependent protein architectures. These observations suggest that ECF distributions may expose modular organisation beyond that recovered by existing domain databases. Receptor-binding proteins – often annotated as tail fibres and tailspikes, are recognised as flagship examples of phage proteins with extensive structural modularity ^11,31,10^. These functions emerged repeatedly in our analyses: tail proteins showed the largest gains in annotation coverage (Figure 3B), tail fibres occupied one of the regions with the highest EF diversity and segregation (Figure 4D), and domain-dark tail fibres were rescued by numerous enriched EFs, including EF0037 (Figure 5B). We therefore focused on a major subtype of receptor-binding proteins, tailspike depolymerases, which are characterised by substantial genetic and functional diversity amid extensive structural modularity ^32^, and asked how ECFs and ECF families (EFs) are organised within those by closely investigating a few cases.

From all rHMMs which (i) were annotated as ‘tail fiber’ or ‘tail spike’ with PHROGs, (ii) contained at least 20 ECF hits, (iii) contained at least two EFs, and (iv) exceeded 600 amino acids in length, we selected five representative depolymerases (`reprseq012341`, `reprseq019028`, `reprseq031110`, `reprseq057378` and `reprseq125186`) whose ECF distributions span a range of ECF–domain relationships (Supplementary Figure S17). Figure 6A maps consensus EFs together with ECOD assignments across these proteins. In all five cases, the dominant pectin lyase-like domain (ECOD 207.2.1) broadly coincided with one or more consensus EFs, despite the latter being derived independently from sequence homology alone. The protein sequence `reprseq019028` additionally contained two annotated domains, a virus attachment domain (ECOD 5092.1.1) and an SGNH hydrolase domain (ECOD 2007.5.1). Whereas the SGNH hydrolase corresponded almost perfectly to EF0012, the virus attachment domain was not recovered by any consensus EF. Thus, ECFs frequently recapitulate conventional domain organisation, while also partitioning proteins in ways that may differ from existing structural classifications.

**Figure 6.**
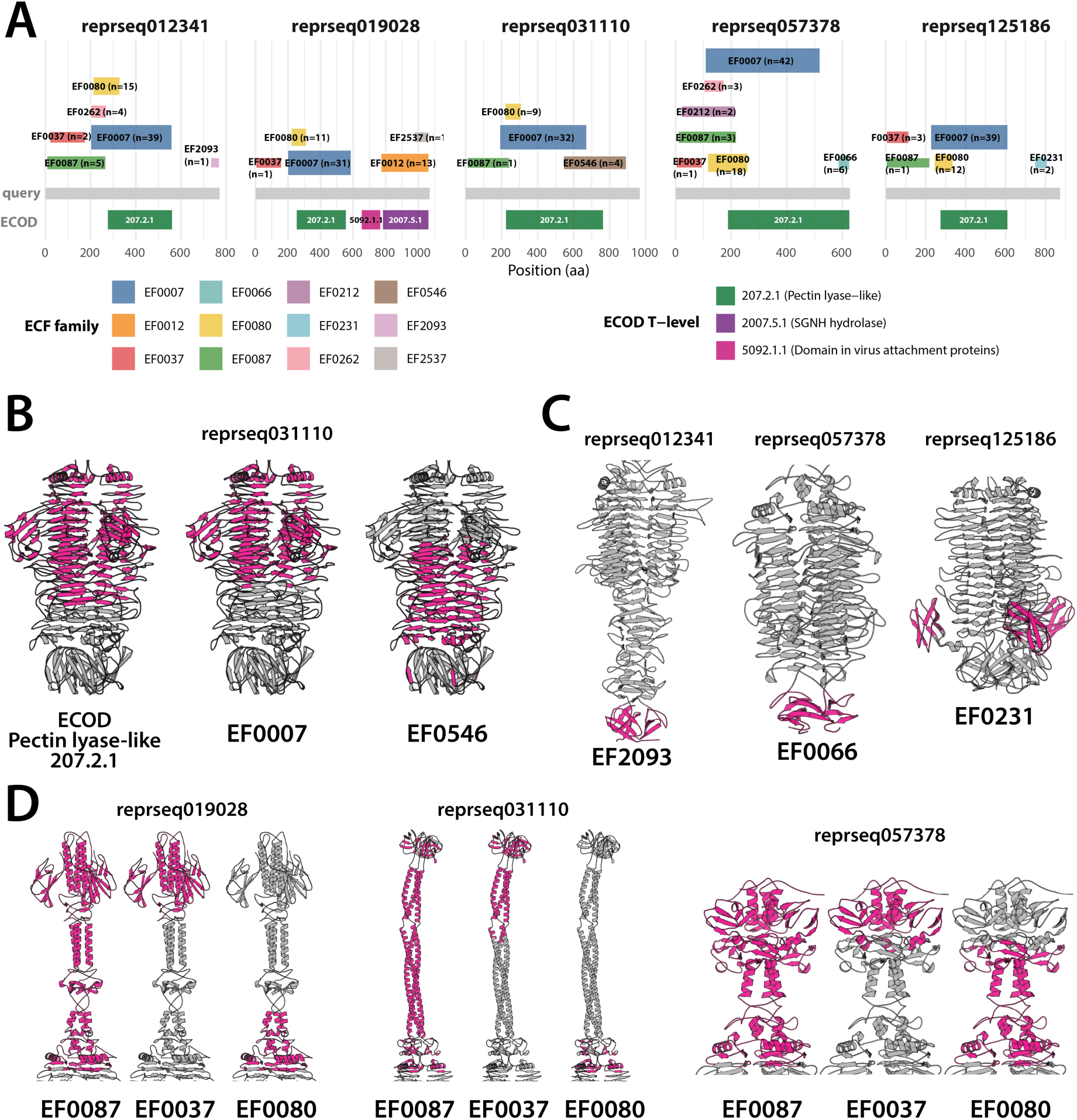
ECF families delineate structural modules in depolymerase-type receptor-binding proteins. **(A)** Consensus ECF family (EF) assignments and ECOD domains mapped onto five representative receptor-binding proteins. Coloured blocks above each grey backbone denote EF families, labelled with the number of contributing ECF hits (*n*); the backbone bar spans the full length of the sequence. ECOD assignments are shown below, coloured and labelled by T-level. Consensus boundaries were derived by quartile-based estimation from the raw hit distributions in Supplementary Figure S17. Three families occurred in all five proteins (EF0007, EF0033, EF0079), whereas EF0013, EF0065, EF0072, EF0212, EF0635, EF2175 and EF2655 were each restricted to a single protein. ECOD assigned a pectin lyase-like domain (207.2.1) to all five proteins and detected no domain outside that region in four of them. **(B)** Two ECF families subdivide a single ECOD domain. AlphaFold3^33^ model of `reprseq031110`, cropped to the C-terminal enzymatic region, with the ECOD 207.2.1 assignment (left) and the EF0007 (centre) and EF0635 (right) assignments highlighted in magenta against grey. EF0007 and EF0635 overlapped the same ECOD domain but resolved it into two offset segments, EF0007 covering the N-proximal coils of the *ω*-helix and EF0635 the C-proximal coils and the region immediately beyond the ECOD boundary. **(C)** ECF families delineate discrete structural elements that received no domain assignment. Cropped models of `reprseq012341`, `reprseq057378` and `reprseq125186`, with EF2175, EF0065 and EF0072 highlighted respectively. Each family corresponded to a compact *ω*-sandwich: distal and C-terminal in `reprseq012341` and `reprseq057378`, and laterally protruding in `reprseq125186`. Neither ECOD nor Pfam assigned any domain to these regions (Supplementary Figure S20). **(D)** Shared N-terminal families occupy structurally dissimilar regions. N-terminal regions of `reprseq019028`, `reprseq031110` and `reprseq057378`, each shown three times with EF0079, EF0033 and EF0080 highlighted in turn. The three families recurred in the same N- to C-terminal order in all three proteins. The regions they occupied differed in fold and in overall geometry, and were modelled at lower confidence than the enzymatic regions (Supplementary Figure S18). All models were predicted with AlphaFold3 as homotrimers and are shown as single chains in (B–D); complete models are shown in Supplementary Figure S18, and every detected family is mapped individually in Supplementary Figure S19.

To investigate regions where ECFs and conventional domains diverged, we predicted AlphaFold3 structures for all five proteins (Supplementary Figure S18) and mapped both consensus EFs (Supplementary Figure S19) and ECOD/Pfam domains (Supplementary Figure S20) onto the resulting models. One striking example occurred in `reprseq031110`, where a single pectin lyase-like ECOD domain was consistently partitioned into two partially overlapping consensus EFs, EF0007 and EF0546 (Figure 6A and 6B). Although structurally continuous, the region behaved at the sequence level as two semi-independent evolutionary modules. Whether these modules correspond to preferential recombination units remains speculative, but their repeated recovery across homologous proteins suggests that they capture biologically meaningful sequence organisation beyond conventional domain boundaries.

Consensus EFs also identified modular units entirely absent from existing domain resources. Figure 6C highlights three examples: EF2093 in `reprseq012341`, EF0066 in `reprseq057378` and EF0231 in `reprseq125186`. None overlapped any ECOD or Pfam annotation, yet each corresponded to a discrete structural element within the AlphaFold3 models, including two C-terminal modules and one insertion motif. Together these examples demonstrate that ECFs recover structurally coherent building blocks invisible to current domain databases. EF0066 and EF0231 also co-occurred in `reprseq094475`, an approximately 3,600-amino-acid receptor-binding protein lacking depolymerase-associated ECFs (Supplementary Figure S21). Although reliable structural modelling of this exceptionally large protein proved infeasible, its predicted monomer architecture suggests that these sequence modules have been reused in a distinct architectural context.

ECFs also revealed conserved modularity where structural correspondence was less obvious. The N termini of all five proteins shared the conserved EF series EF0087–EF0037–EF0080 (Figure 6A). We focused on three of those (`reprseq019028`, `reprseq031110` and `reprseq057378`) because their AlphaFold3 models were sufficiently well resolved to permit structural comparison in this region. Despite their shared sequence architecture, the corresponding models exhibited limited structural similarity, particularly within elongated *ω*-helical coiled coil regions predicted with comparatively low confidence. This discrepancy may therefore reflect limitations of current structure prediction, genuine structural divergence despite shared ancestry, or both. Regardless of its origin, the reproducible recovery of homologous EFs across these proteins indicates that conserved sequence modules persist even where structural relationships are ambiguous. Together, these examples demonstrate that ECFs complement rather than replace conventional domain annotation, providing an additional sequence-derived layer of modular organisation that can subdivide known domains, identify previously unrecognised structural motifs, and detect homologous modules that remain difficult to recognise using structural information alone.

## Discussion

This study provides a resource catalogue of 13,035 Evolutionarily Conserved Fragments (ECFs), grouped into 3,167 families, from 5,396 RefSeq bacteriophage genomes. ECFs were defined from sequence alone and without pre-imposed domain boundaries, but exploited the sensitivity of HMM–HMM comparisons to identify homology signal too distant for sequence searches to recover. That signal correlated with existing domain resources (ECOD and Pfam), but ECFs also differed from them: their length distribution was more extreme at both ends, and their organisation was inherently nested, with shorter insets recurring inside longer containers. Direct comparison resolved a heterogeneous spectrum of relationships to established domains, from subdomains through equivalents to superdomains, though the largest class had no counterpart in either resource. Consequently, ECFs annotated 50% of representative proteins, in contrast to 25% for Pfam and 20% for ECOD, and identified 23,925 domain-dark proteins that lacked any Pfam or ECOD hit. Among the 2,959 of these with a confident functional annotation, 242 EF–function associations were significant, indicating that EFs can act as functional signatures where domain databases are silent, with the strongest signal in structural tail proteins. Because many ECFs overlapped into ECF Families (EFs), protein architecture could be described along two largely independent axes, EF diversity and EF segregation, which varied with function and exposed modularity that domain annotation left implicit. Examined closely in tailspike depolymerases, EFs subdivided a single domain, delineated unannotated structural elements, and recurred where structural similarity was weak. None of these patterns could be inferred from domain or structural annotation alone.

The defining feature of ECFs is not simply that they are derived from sequence, but that they partition proteins according to evolutionary conservation rather than imposing structural boundaries. By asking which regions recur across otherwise distinct protein families, ECFs aim to recover homologous building blocks: units that have been retained, exchanged or recombined over evolutionary time. This places them in the tradition of *de novo* domain-detection approaches such as ADDA ^24^ or EVEREST^22^, but also exposes the limitation of describing all such units as domains. Although many ECFs correspond closely to established domains, conservation need not respect the boundaries of an independently folding unit. Domains that remain tightly linked during evolution may emerge as composite superdomains, whereas ancient motifs or highly conserved regions shared amongst domains may emerge as subdomain fragments ^34^. ECFs therefore provide an evolutionary decomposition of protein sequence that is related to, but fundamentally distinct from, its structural decomposition. Their value is consequently greatest not as an alternative to structural annotation, but as a complementary reference frame for asking whether what folds together also evolves together. In our demonstrated example of tailspike depolymerases, combining ECFs with domain assignments and structural models revealed precisely this distinction: ECFs identified conserved units within an independently folding receptor-binding domain, exposed mosaicism below the classical single-stranded *ε*-helix domain level, and connected regions whose structural similarity alone was weak. Such evolutionary building blocks may ultimately provide a more informative vocabulary for understanding—and potentially engineering—the modular assembly of phage proteins.

A broader way to view ECFs is as a protein-scale analogue of ancestry-painting approaches in population genetics. Methods such as fineSTRUCTURE ^35^ or ADMIXTURE ^36^ in eukaryotes, and BRAT^37^ or fastGEAR ^38^ in bacteria, describe genomic DNA by assigning its successive regions to the reference populations with which they share the greatest affinity, thereby revealing a mosaic of local evolutionary histories. ECFs apply a similar representation to proteins, mapping different regions of a sequence to conserved fragment families sampled across phage sequence space. The analogy only holds partially as proteins are far shorter than genomes, and a single region may match several overlapping ECFs rather than one discrete ancestral population. Yet these patterns remain informative. Overlapping ECFs indicate locally related evolutionary signals, whereas spatially segregated ECFs reveal blocks with different affinities. Phage proteins can therefore be viewed not only as arrangements of structural domains, but as mosaics of fragments with partially separable evolutionary histories. Such painting is particularly useful for exceptionally long, poorly annotated proteins, such as tape-measure proteins, for which ECF-based “painting” describes an internal composition that domain databases do not capture: in our data, these proteins carried repeated, spatially segregated ECF families, consistent with a modular, repeat-based architecture.

A potential limitation of ECFs is that their mappings can appear considerably noisier than those produced by manually curated resources such as Pfam or ECOD. Individual proteins may be covered by several overlapping ECFs, and some regions do not resolve into a single, sharply bounded unit. The redundancy is partly a consequence of design. ECF construction deliberately minimises manual consolidation in order to preserve the structure present in the underlying sequence relationships. The resulting redundancy and nesting are therefore not merely technical artefacts: shorter ECFs frequently occur within broader ones and can connect otherwise dissimilar regions of protein space, revealing conserved links that would be lost if related profiles were collapsed into a single curated family. Conversely, regions that produce diffuse or inconsistent ECF patterns may reflect sequences for which evolutionary conservation is itself poorly localised, including disordered, rapidly evolving or elongated *ω*-helical regions in long tail fibres. Such patterns should not be interpreted uncritically, but neither should they be treated simply as failed annotation. ECFs are consequently most informative when the full mapping pattern is considered—including overlap, nesting and the absence of clear boundaries—rather than when each hit is treated as an independent, definitive domain assignment.

A few important limitations need to be acknowledged. ECFs were only as representative as the protein space from which they were built. The catalogue derived entirely from RefSeq phage genomes, which represent a biased and incomplete sample of global phage diversity — enriched for well-studied tailed phages of cultivable bacterial hosts and depleted for environmental lineages. Application to an independent metagenomic dataset confirmed that ECFs extended coverage beyond what Pfam and ECOD achieved, but also revealed a proportionally larger unannotated fraction than in RefSeq, consistent with substantial missing diversity. ECF construction also depended on fragment frequency: rare fragments present in few protein families were less likely to survive the clustering and coverage filters, and unequal sampling of phage lineages in RefSeq propagated into unequal ECF representation. Finally, ECFs were sequence-defined and not structurally validated at scale. The receptor-binding protein case study confirmed structural coherence for four proteins, yet that correspondence remained untested across the full catalogue. Systematic structural validation using AlphaFold or experimental structures would strengthen the biological interpretation of ECF family boundaries considerably.

A complete account of protein evolution will require identifying not only what folds independently, but what is inherited, exchanged and recombined as a unit. Our results show that these evolutionary units do not always coincide with classical domains, particularly in the highly modular and poorly annotated proteins of phages. By revealing conserved sequence building blocks across this space, ECFs provide a first step towards reconstructing how complex phage proteins are assembled—and towards using that knowledge to predict and design their functions.

## Methods

### Data collection and rHMM construction

We downloaded all (*n* = 5, 396) NCBI RefSeq genomes from the INPHARED database (Dec 2024 release)^39^. We then ran prodigal-gv ^40^ to predict coding sequences, removing any sequence with more than 5 unknown characters, yielding 540,772 protein sequences. All protein sequences were clustered with mmseqs2 (release 17-b804f) ^41^ using the following parameters: minimum sequence identity = 0.3, sensitivity = 7.5, coverage = 0.95 (otherwise default), resulting in 148,010 clusters. A representative sequence was then extracted from each cluster, and a multiple sequence alignment was built following the HHpred pipeline ^30^, namely by aligning each representative protein sequence against the UniClust30 database (2023_02 release) ^42^ using hhblits from the hhsuite package (release 3.3.0) ^43^ and preserving up to 200 most diverse representatives, yielding 148,010 representative rHMMs.

### Function and domain assignment in rHMMs

All rHMMs were searched against four external databases using hhsearch: PHROGs (v4)^44^, ECOD (version 20230309) ^13^, Pfam (v37) ^45^ and an antidefence protein database ^46^ compiled in ^10^. Functional annotation was based on PHROGs and the antidefence database; domain annotation was based on ECOD and Pfam. For PHROG-based annotation, hits were retained if they met a minimum probability of 95%, minimum alignment length of 30 amino acids, and minimum query and subject coverage of 60% each. For antidefence annotation, the search was conducted with the antidefence sequences as query and rHMMs as subjects; the same probability, length and coverage thresholds were applied, with query and subject roles reversed accordingly. Hits passing these filters were ranked by a composite score (query coverage × subject coverage), then by percentage identity and e-value. Each rHMM was assigned the single top-ranking annotation (PHROG or antidefence). A function was retained as a primary annotation only if the corresponding function contained at least 500 sequences and at least 100 rHMMs carried that annotation; functions not meeting these criteria were excluded from downstream functional analyses unless stated otherwise. Annotations corresponding to non-specific terms (“tail”, “structural protein”) or “unknown” were also excluded. For domain annotation, rHMM hits to ECOD and Pfam were filtered at a minimum probability of 70%, maximum e-value of 0.1, and minimum hit length of 20 amino acids, applying additional filters when necessary (see Results).

### Detection of Query Evolutionarily Conserved Fragments (qECFs)

Using the all-by-all comparison of rHMMs described above, for each rHMM, all hits to a given query were clustered according to the start (qstart) and stop (qstop) position of the hit in the query, excluding self-hits and hits with probability less than 90%. This filtering resulted in 118,377 rHMMs that passed criteria and were considered in the clustering step. Clustering was carried out using DBSCAN with epsilon=0.5 and minPts = 5, implemented in the sci-kit learn Python package ^47^. The purpose of this clustering was to find candidates for query-ECFs (qECFs), which can be thought of as projections of true ECFs onto each query (rHMM).

The value of epsilon was chosen to strike a balance between producing concise, meaningful clusters and effectively managing noise, ensuring an optimal distinction between linking only closely related sequences into a qECF (low epsilon) and the over-merging of distinct sequences into a qECF (high epsilon). On the other hand, the choice of minPts safeguards against the dilution of signal from rare proteins while minimising the influence of uncertain hits. This approach resulted in 176,122 qECF candidates.

For each qECF candidate, we calculated two important statistics: average hit overlap (AHO) and average coverage (AC). AHO was obtained as the mean overlap between all pairs of hits within the candidate qECF, where an overlap was defined as the length of hit intersection divided by the length of the hit union, each determined from the coordinates in the query. To calculate the average coverage (AC) for each candidate qECF, we first established the boundaries of the qECF. We set the start boundary at the first quartile (25th percentile) of all start positions of hits within the qECF, and the end boundary at the third quartile (75th percentile) of all end positions. This approach was chosen to minimise the impact of noise in the distribution of start and end positions, effectively removing the outlying, less reliable flanks of the qECF. The length between the qECF start and end was defined as the qECF span. AC was then calculated as the mean hit coverage, defined as the average proportion of the qECF span covered by each hit.

To determine the AHO and AC value thresholds which constitute a high-quality qECF, we considered only rHMM hits to the Pfam database with a minimum probability of 95% and subject coverage of 90%. For every candidate qECF, we then calculated a maximum overlap between its span and the Pfam hits within the rHMM and considered only those qECFs for which such an overlap was at least 80%. The qECFs fulfilling these criteria were considered as Pfam-equivalent. The cut-off AHO and AC values were determined by truncating the lower 5% of the distribution, effectively removing the lowest values to minimise the impact of outliers. The following cut-off values were obtained: AHO = 0.81 and AC = 0.87, which were used to filter low quality qECFs. Finally, all qECFs with the length shorter than 20 amino-acids were also excluded to avoid the inclusion of peptides, thus retaining 143,955 qECFs.

### Construction of qECF alignments

For each qECF, a multiple sequence alignment (MSA) was created from the alignments yielded by the all- vs-all search of all rHMMs. The qECF alignments were built from the rHMM (query) sequence between the denoised boundaries (start and end positions) and sequences of hits (subjects) associated with the corresponding qECF cluster. Importantly, each hit was verified before being added to the MSA: hits with e-value higher than 10^−2^ or percentage identity lower than 10% were excluded. If there were fewer than 4 hits that met these criteria (hence 5 sequences in the MSA), the MSA was not created and a given qECF was excluded from further analyses. The approach resulted in a final set of 121,594 qECFs.

### Construction of ECFs from qECFs

For each qECF, we constructed HMM profiles from their respective MSAs and performed an exhaustive all-vs-all comparison via hhsearch retaining only hits with the minimum probability of 95%. Then, for each qECF pair, denoted A and B, we determined query and subject coverage values (qcov and scov) in the following way. Query coverage was the minimum of: qcov when A served as the query and scov when A served as the subject; subject coverage was the minimum of: scov when A served as the query and qcov when B served as the subject. Similarly, for each pair, the probability and percentage identity were conservatively derived as the lesser of the two alignment evaluations (A vs B, and B vs A). In instances where one alignment was absent, the available value was used. All pairs of qECFs were then considered as overlapping when their coverage values (qcov and scov) were both at least 90%. Only overlapping pairs were considered in the next step. A weighted network was then created for all pairs with the weight given as qcov×scov (given qcov≥ 0.5 and scov≥ 0.5) and provided as input into MCL with inflation factor of -I 1.3. This resulted in 13,035 clusters, which we considered as putative ECFs.

### Construction of ECF Alignments

For each resulting MCL cluster, we reconstructed a qECF similarity graph where nodes corresponded to qECFs in the cluster and edges corresponded to previously linked qECFs (i.e., coverage of minimum 90%). We then identified a qECF with the highest node centrality in this network (i.e., linking the highest proportion of qECFs), which then served as the starting alignment. The ECF alignment was then prepared with hhalign by collecting MSAs of all other qECFs and successively aligning them against the starting alignment with default values. The final alignment was then processed with `hhfilter` with maximum identity (`-id`) of 95% and minimum coverage (`-cov`) of 50%, resulting in 13,035 ECF alignments which were converted into a hhsuite database.

### Mapping ECFs to domain databases and rHMMs

ECF profiles were searched against ECOD and Pfam, and all rHMMs were searched against the ECF database, using hhsearch. For analyses of ECF content in rHMMs (annotation coverage, domain-dark proteins, architecture), hits were further restricted to probability ≥ 95%, e-value ≤ 0.1, hit length ≥ 20 aa and ECF (subject) coverage ≥ 70%. To support sensitivity analyses, a second set of ECF-to-ECOD and ECF-to-Pfam hit tables was produced using a lenient e-value threshold of 10, with no other filters applied (see Results). For each ECF pair, conservative query and subject coverage values were computed from hits with probability ≥ 95%: qcov was taken as the minimum of qcov(A→B) and scov(B→A), and scov as the minimum of scov(A→B) and qcov(B→A).

### Construction of ECF families (EFs)

ECF families were constructed by graph-based clustering of the all-vs-all ECF comparison. An undirected weighted graph was built in which nodes represented ECFs and edges connected pairs with mutual query and subject coverage of at least 50% and alignment probability of at least 95%. Edge weights were set to the product of qcov, scov, and probability. The graph was clustered using MCL with an inflation factor of 1.3, chosen by visual inspection of the resulting clique structure and the functional coherence of ECFs within and between clusters. The 3,167 resulting clusters were designated ECF families (EFs).

### Extraction of metagenome-derived proteins

To evaluate ECF annotation coverage in a dataset independent of the RefSeq training set, we sampled 2,000 phage genomes from the PhageScope database ^48^. PhageScope clusters were first screened to remove any cluster containing a GenBank or RefSeq genome; this ensured that selected genomes were not represented in the rHMM dataset. From the remaining clusters, 2,000 were drawn at random. Within each cluster, the single best representative genome was selected by prioritising assembly completeness, then genome length. All 102,676 proteins from the 2,000 representative genomes were retained for analysis. Protein sequences were then searched against ECF, ECOD, and Pfam HMM databases using hhsearch, applying the same filtering thresholds as for rHMMs: minimum probability of 70%, maximum e-value of 0.1, and minimum hit length of 20 amino acids. The fraction of proteins receiving at least one hit from each database was computed per functional category and compared across databases.

### ECF–domain relationship classification

Each ECF was assigned to one of six relationship classes relative to ECOD, based on the alignment probability, e-value, query coverage (qcov), and subject coverage (scov) of its best ECOD hit. Assignments were made in priority order as follows. An ECF was classified as *equivalent* to an ECOD domain if probability ≥ 95%, e-value ≤ 0.01, qcov ≥ 0.8, and scov ≥ 0.8, indicating reciprocal near-complete coverage. An ECF was classified as a *subdomain* of an ECOD domain if probability ≥ 95%, e-value ≤ 0.01, qcov ≥ 0.9, and scov ≤ 0.6, indicating that the ECF is largely contained within a larger ECOD domain. An ECF was classified as a *superdomain* if probability ≥ 95%, e-value ≤ 0.001, qcov ≤ 0.6, and scov ≥ 0.9, indicating that the ECF spans beyond a single ECOD domain. Remaining ECFs with confident hits (probability ≥ 95%) that did not meet any of the above criteria were classified as *related*. ECFs that retained at least one ECOD hit passing the search thresholds but no hit reaching the confident probability threshold were classified as *distant*. ECFs with no ECOD hit meeting the minimum thresholds (e-value of 0.1) were designated as *new* and treated as annotation-orphans for the purposes of functional enrichment analysis. The same classification scheme was applied independently to Pfam hits for comparison. For each ECF with at least one confident ECOD hit, we counted the number of unique ECOD X-groups (the highest hierarchical level in the ECOD classification) spanned by its hits. This quantity was used to assess whether individual ECFs link multiple distinct structural lineages.

### Protein architecture metrics

To characterise how ECF families organise within individual proteins, we computed a suite of per-rHMM architecture metrics from ECF-to-rHMM hit tables. All metrics were derived from hits filtered at probability ≥ 95%, e-value ≤ 0.1, minimum hit length ≥ 20 aa, and ECF (subject) coverage ≥ 70% — the same thresholds used for annotation coverage analyses. Each hit was assigned to the ECF family of its corresponding ECF.

#### Positional consistency

To assess the positional consistency of repeated occurrences of the same ECF family within a rHMM, we computed the *within-family average hit overlap* (AHO), in similarity to the qECF metric introduced earlier. For each rHMM–ECF family combination where the family contributed at least three hits, all pairwise Jaccard overlaps were computed between hit intervals (intersection length divided by union length), and the mean was taken as the within-family AHO for that rHMM–ECF family pair.

#### Coverage

Each hit was expanded to its per-residue footprint (all positions from `qstart` to `qend`), and the *coverage fraction* (`coverage.fraction`) was defined as the number of positions covered by at least one ECF hit divided by the rHMM length. We also counted the *total number of ECFs* (`n.ecf.hits`) found in the rHMM.

#### Compositional complexity

The *number of ECF families* (`n.fams`) was the count of distinct ECF families detected across all hits to a given rHMM. To quantify positional family diversity while accounting for unequal hit multiplicity, we computed a Simpson-based effective number of families at each covered position. For each position, the proportion of hits belonging to family *f* was *p_f_* = *N_f_*/Σ*_f_ N_f_*, where *N_f_* was the number of hits from family *f* at that position, and the effective number of families was 1/Σ*_f_p_f_*^2^. The *mean eflective number of families* (`mean.eff.fam`) was the average of this quantity across all covered positions.

To capture the spatial patchwork of family dominance along the protein, we computed the *number of dominant-family segments* (`n.dom.segments`). Each covered position was assigned the family with the highest hit multiplicity at that position, breaking ties alphabetically. Positions where the dominant family’s share fell below 0.6 were marked as ambiguous and excluded. The number of dominant-family segments was then counted as the number of contiguous runs among covered positions (uncovered positions did not break a run).

#### Spatial segregation

We defined two complementary segregation indices to measure the degree to which ECF families partition into spatially distinct blocks along the protein. Both were defined only for proteins carrying two or more families. The *overlap-based segregation index* (`segregation.overlap`) measured range separation. For each protein–family pair, the positional range was [min(pos), max(pos)] across all covered positions. For each pair of families within a protein, overlap was computed as the length of the intersection of their ranges divided by the length of the shorter range, yielding a value between 0 and 1. The index was defined as 1 ↑ *O*, where *O* was the mean pairwise overlap fraction. The *barycentric segregation index* (`segregation.bary`) instead measured the separation of family mass centres. For each protein–family pair, the barycenter was *µ* = Σ(pos × *N*)*/* Σ *N*, where *N* was the hit multiplicity at each position. For each pair of families, the normalised distance was |*µ_i_ – µ_j_|/L*, where *L* was the rHMM length. The barycentric segregation index was the mean pairwise normalised distance.

## Data Availability

All data supporting this study are publicly available. Input genomes were obtained from the INPHARED database (December 2024 release) ^39^, and the 2,000 phage genomes used for independent validation were sampled from PhageScope ^48^. Annotation resources are publicly available: PHROGs v4, ECOD (version 20230309), Pfam v37, UniClust30 (2023_02) and the antidefence protein database of Samuel et al. ^46^. The ECF database (HH-suite format, comprising the ECF alignments and HMM profiles), representative sequences and rHMMs, annotations, hit tables, ECF and ECF family assignments, protein architecture metrics and the PhageScope-derived tables are available at FigShare (https://doi.org/10.6084/m9.figshare.34887162). The ECF database is also provided as downloadable files (with SHA-256 checksums) attached to release v1.0.2 of the analysis repository (https://github.com/rmostowy/ECFs/releases/tag/v1.0.2).

## Code Availability

The workflow for constructing ECFs, from genome processing and representative sequence clustering through rHMM construction and all-versus-all comparison to ECF and ECF family generation, is available at flow: `notebooks/ECF-Workflow-final-v25082025.ipynb`) and (main workflow `notebooks/ECF-Workflow-final-v25082025.ipynb`) and https://github.com/bioinf-mcb/phage-protein-modularity-data (branch `ecf-paper-2026`). The code for annotation, domain and ECF analyses, protein architecture metrics and figure generation is available at https://github.com/rmostowy/ECFs, which reads the outputs of the workflow above. Version v1.0.2 of the analysis code used in this study is archived on Zenodo (https://doi.org/10.5281/zenodo.23953341).

## Author contributions

The following contributions can be distinguished by different authors based on the CRediT (Contributor Roles Taxonomy) system:

- **Conceptualization**: SDH, RJM
- **Methodology**: BS, KS, RJM
- **Software**: KS, JL, RJM
- **Validation**: BS, KS, RJM
- **Formal Analysis**: BS, KS, JL, RJM
- **Investigation**: BS, KS, SDH, RJM
- **Resources**: RJM
- **Data Curation**: BS, KS, RJM
- **Writing – Original Draft**: BS, KS, RJM
- **Writing – Review & Editing**: BS, KS, SDH, RJM
- **Visualization**: BS, RJM
- **Supervision**: RJM
- **Project Administration**: RJM
- **Funding Acquisition**: RJM

