## Supplementary Figures for "Evolutionarily Conserved Fragments reveal modular architecture and expand functional coverage in bacteriophage proteins"

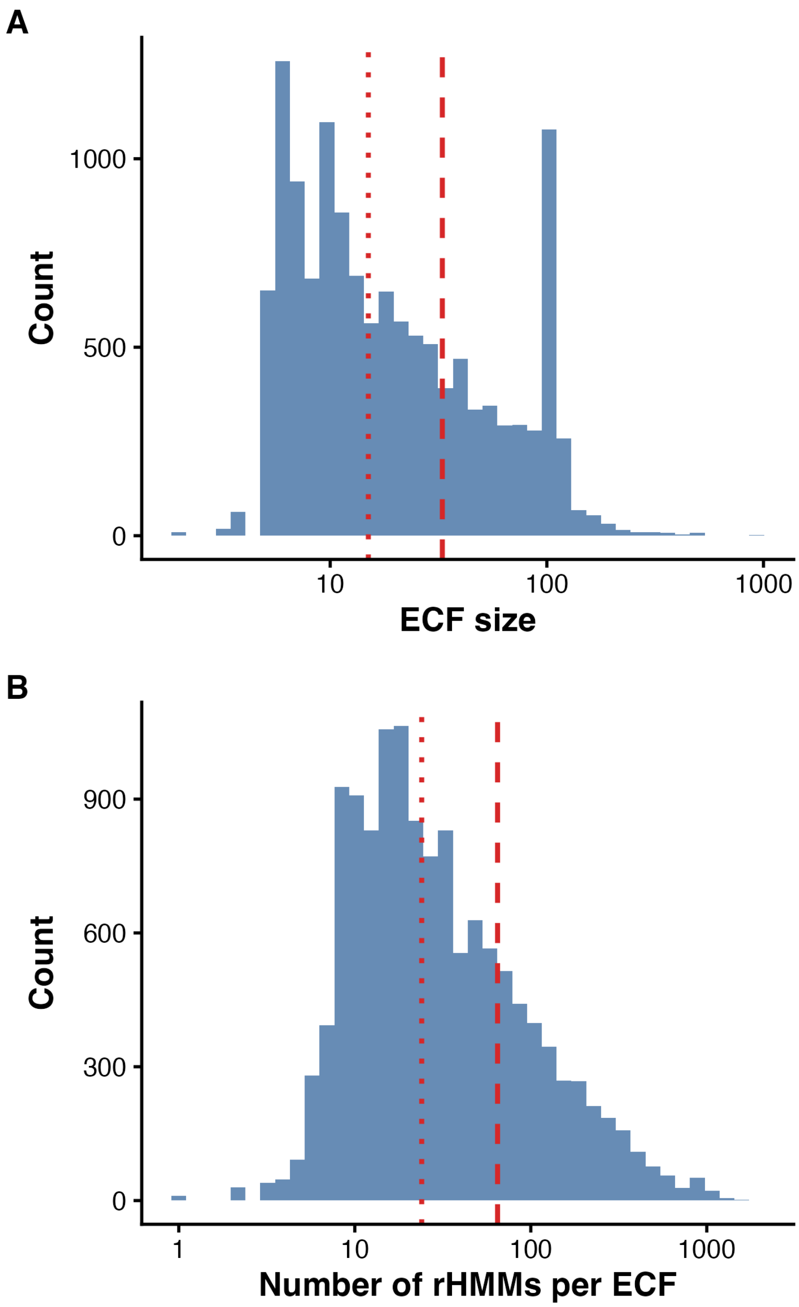


**Figure S1. ECF size distribution**. (A) Distribution of ECF alignment size, measured as the number of sequences in each ECF multiple sequence alignment. Dashed and dotted red lines indicate the mean and median, respectively. (B) Distribution of the number of rHMMs detected per ECF using the following additional filtering criteria: probability $\geqslant$95%, qcov$\geqslant$0, scov$\geqslant$0.7. Both distributions are right-skewed on a log scale, indicating that most ECFs are modest in size and reuse frequency, while a minority are substantially larger or more broadly distributed.

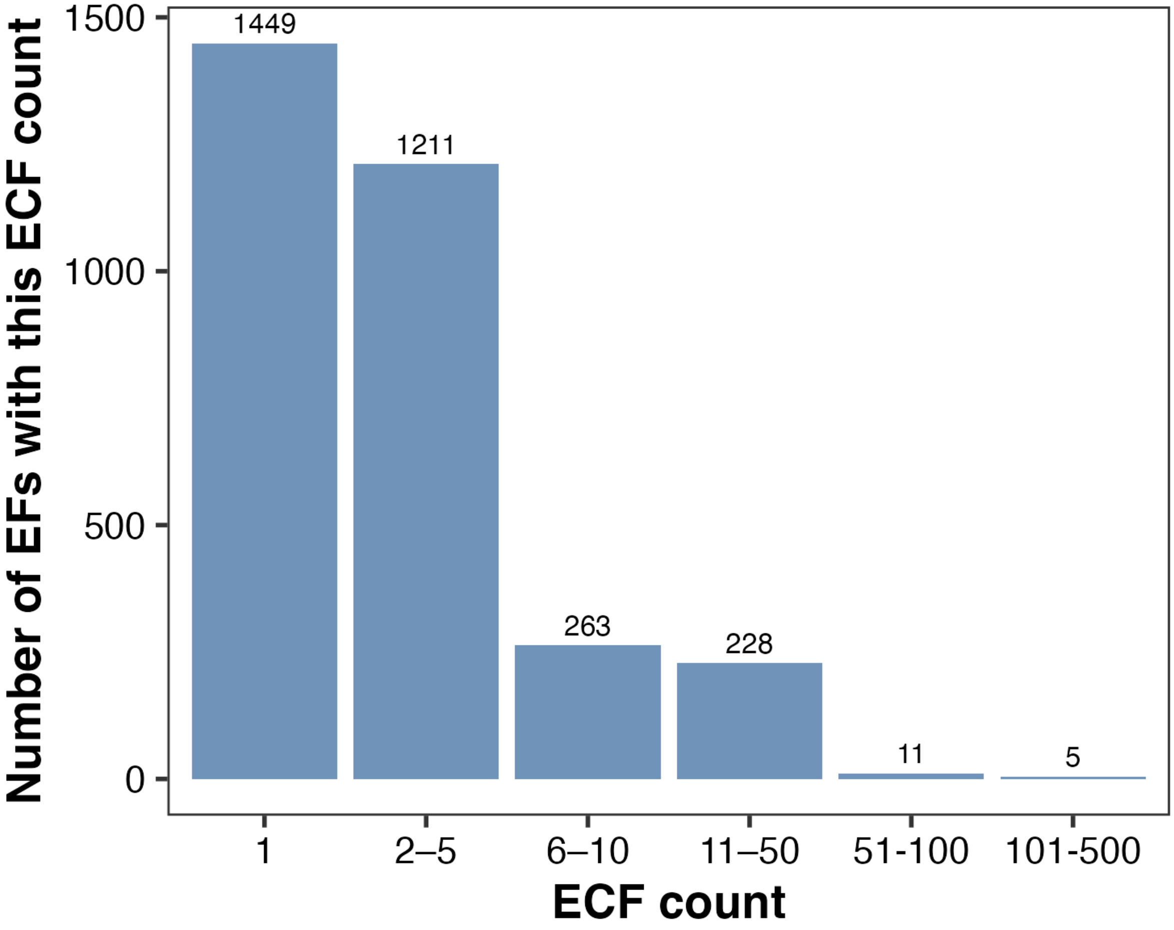


**Figure S2. ECF count per ECF family (EF).** Number of ECF families (EFs) containing a given number of constituent ECFs. The majority of families contain one (1,539) or two to five (1,252) ECFs, with very few families (15) containing more than fifty ECFs, reflecting the broad diversity of conserved fragments across phage protein space.

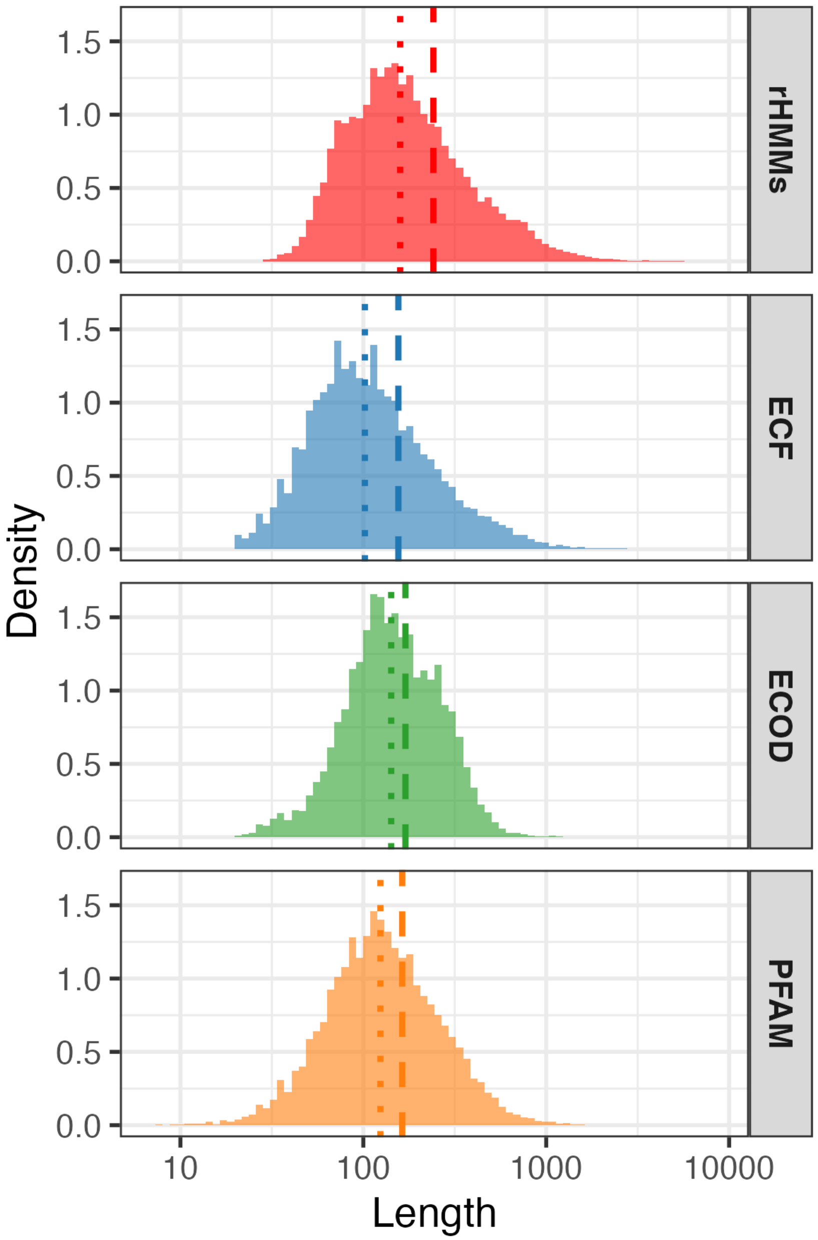


**Figure S3. Length distributions of rHMMs, ECFs and domain-database entries**. Length distributions of rHMMs, ECFs, ECOD domains and Pfam domains on a log scale. Dashed and dotted vertical lines indicate the mean and median for each set, respectively.

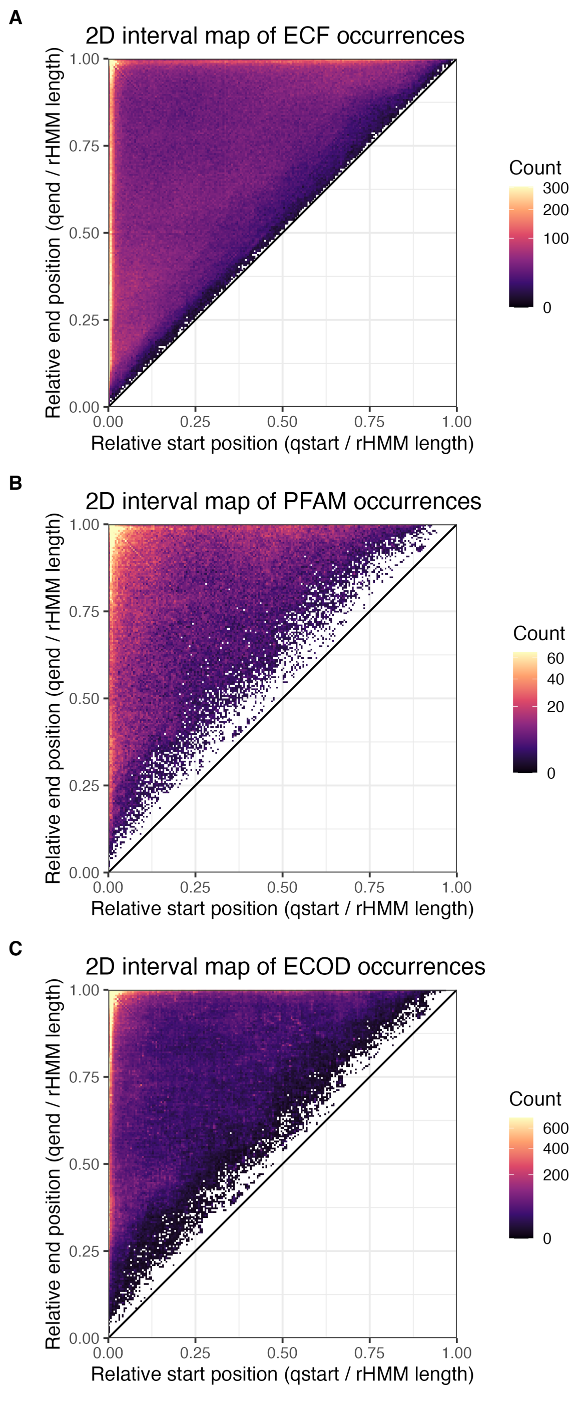


**Figure S4. Positional distributions of ECF, Pfam and ECOD occurrences within rHMMs.**. Two-dimensional interval maps showing the normalised start and end positions of (A) ECF, (B) Pfam, and (C) ECOD hits within rHMMs. Each point represents a single hit, with relative start (qstart / protein length) on the x-axis and relative end (qend / protein length) on the y-axis. All three databases show qualitatively similar positional patterning, with enrichment along the diagonal and at terminal regions, though ECF hits are quantitatively denser, consistent with the shorter and more numerous nature of ECF fragments. The following additional filtering criteria were used to detect ECFs/domains in rHMMs: probability $\geqslant$95%, qcov$\geqslant$0, scov$\geqslant$0.7.

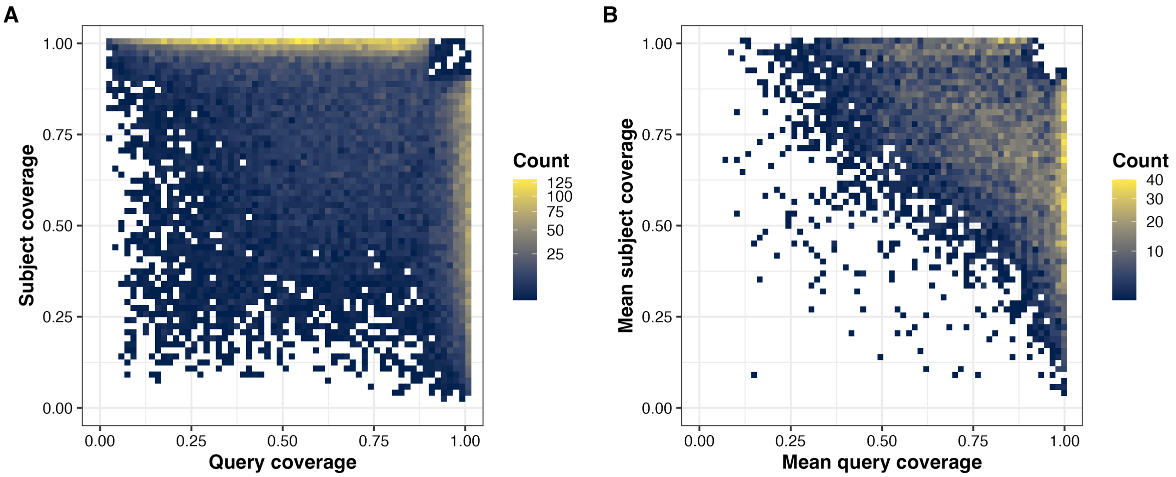


**Figure S5. Coverage asymmetry of ECF–ECF alignments at the alignment and at the ECF level.** (A) Query against subject coverage for all individual high-confidence ECF–ECF alignments (probability $\geqslant$ 99%), shown as a two-dimensional binned density with a $\log_{10}$ count scale. (B) The same quantities summarised per ECF as the mean query and mean subject coverage across all of its connections. Both panels reproduce the L-shaped distribution seen for per-ECF medians in Figure 2A, indicating that coverage asymmetry was a general property of the catalogue rather than an artefact of either the summary statistic used or a small number of highly connected ECFs.

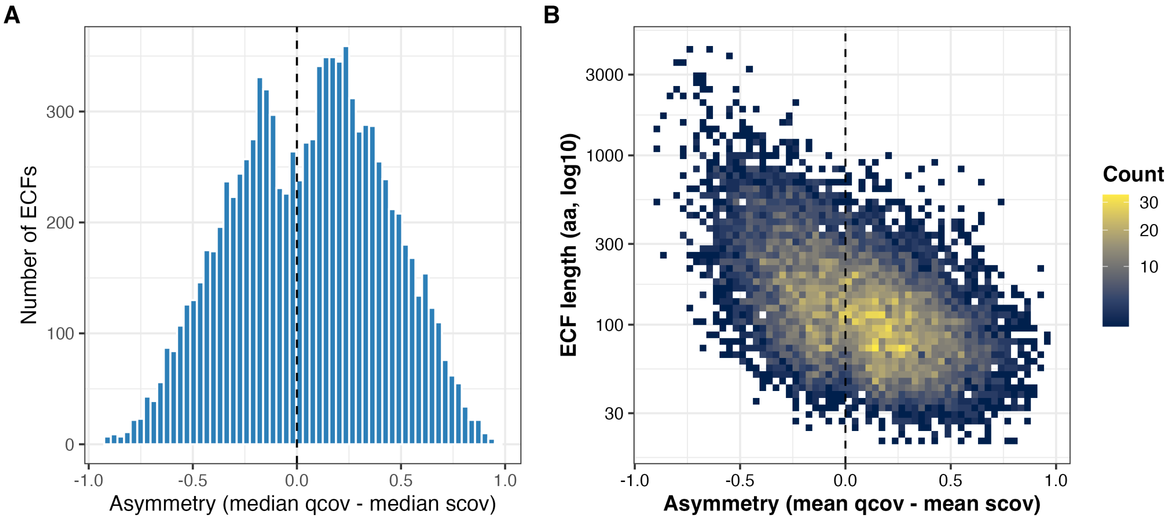


**Figure S6. Per-ECF coverage asymmetry and its relationship to ECF length.** (A) Histogram of per-ECF asymmetry, defined as median `qcov` minus median `scov` across all high-confidence connections (probability $\geqslant$ 99%). Positive values identify ECFs predominantly contained within larger ones (insets); negative values identify ECFs that predominantly act as containers. The distribution spanned the full range from $-1$ to $+1$ and was shifted slightly positive, indicating that insets were modestly more frequent than containers across the catalogue. The dashed line marks zero asymmetry. (B) Binned density of ECF length (amino acids, $\log_{10}$ scale) against mean coverage asymmetry, with a $\log_{10}$ count colour scale. Inset behaviour concentrated among shorter ECFs and container behaviour among longer ones. The dashed line marks zero asymmetry.

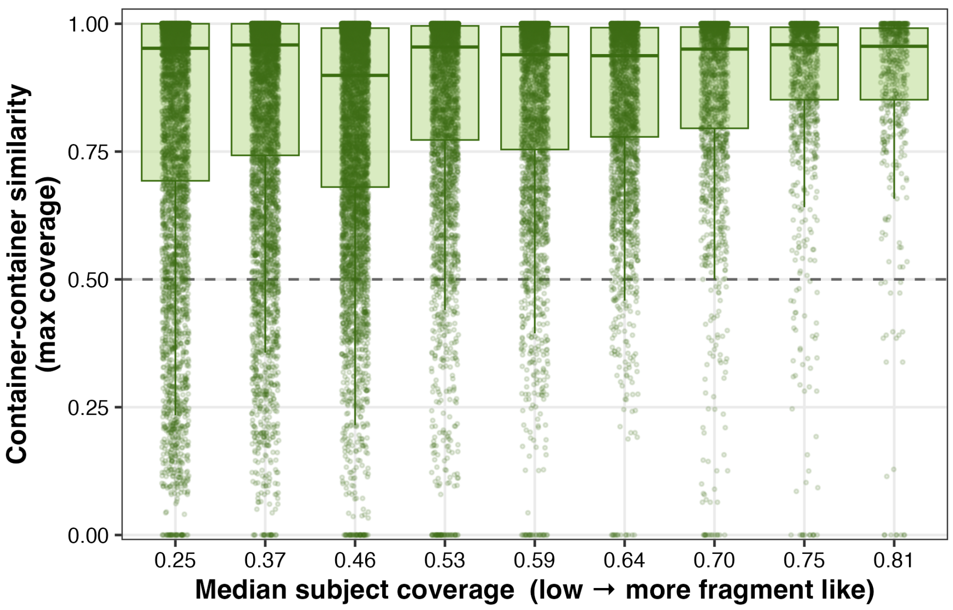


**Figure S7. Full distribution of container–container similarity across degrees of containment.** For each inset ECF, every pair of ECFs containing it was scored by the maximum pairwise coverage between the two containers. Distributions are shown for deciles of the median subject coverage of the inset ECF, plotted left to right from stronger to weaker containment; x-axis labels give the decile midpoints. Boxes span the interquartile range with the median marked, individual container pairs are overlaid as jittered points, and the dashed line marks the 0.5 threshold below which a pair was treated as unrelated. Figure 2B collapses each decile to the proportion of pairs falling below that threshold; the full distributions are shown here to make explicit that near-zero container–container similarity occurred across the whole range of containment rather than only at one extreme.

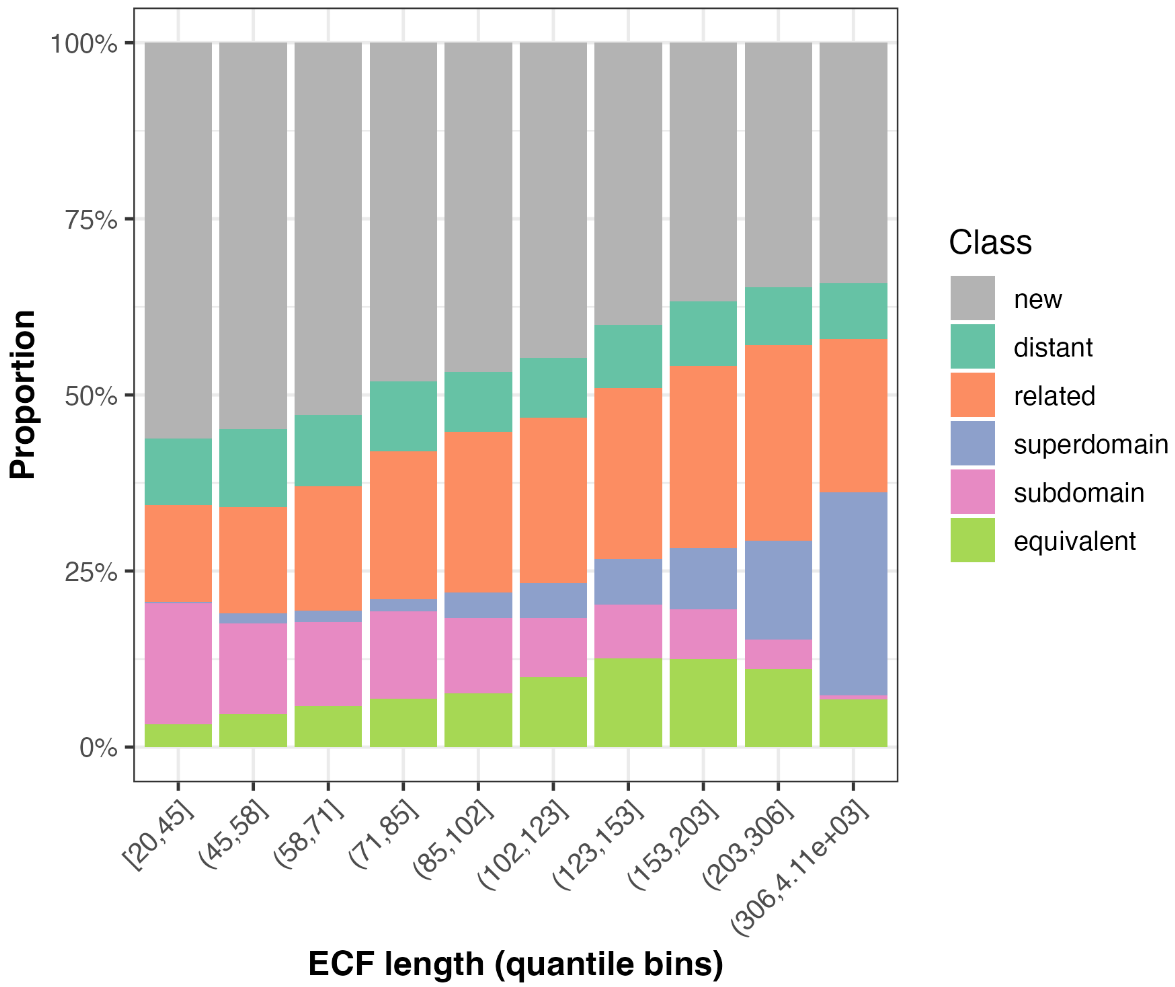


**Figure S8. ECF–Pfam relationship class distribution across ECF length deciles.** Proportional distribution of ECF–Pfam relationship classes across ECF length deciles, using the same classification criteria as in Figure 2C applied to Pfam hits.

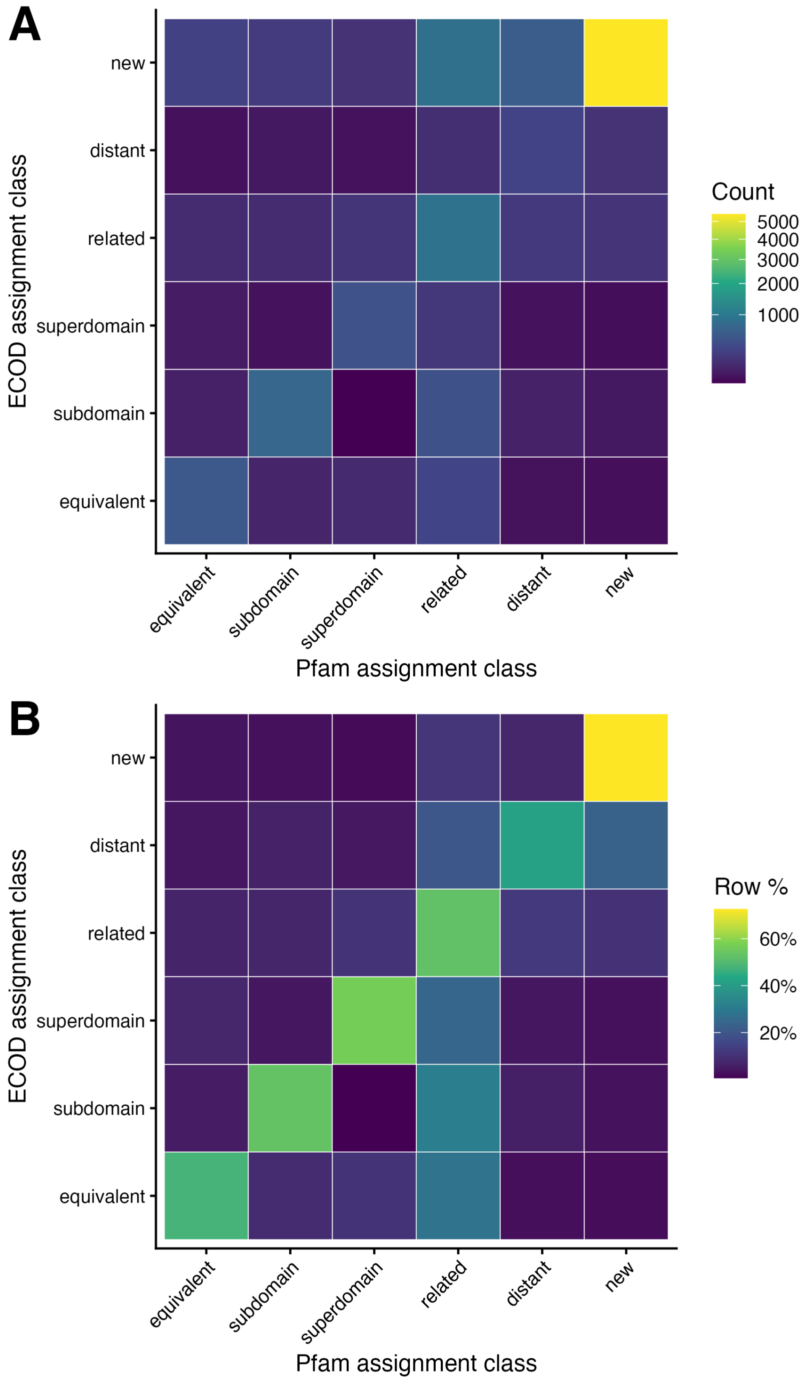


**Figure S9. Agreement between ECOD- and Pfam-based ECF relationship class assignments.** (A) Raw counts and (B) row-normalised proportions of ECF class assignments in ECOD (rows) and Pfam (columns), including an explicit absent class for ECFs lacking any hit in a given database. A Pearson chi-squared test confirmed a significant association between the two classification schemes ($\chi^2(25) = 5757$, $p < 2.2 \times 10^{-16}$; Cramér's $V = 0.391$).

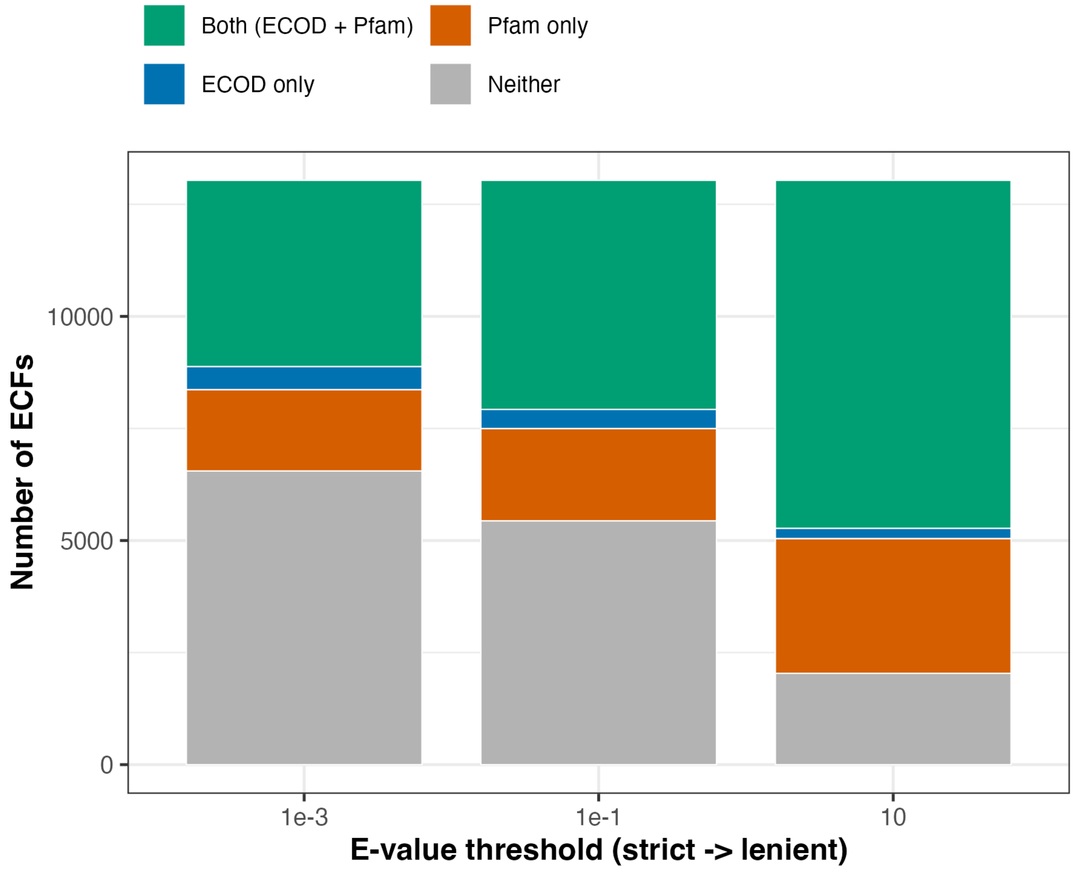


**Figure S10. ECF coverage by ECOD and Pfam across three detection thresholds.** Number of ECFs with a hit to ECOD only, to Pfam only, to both databases or to neither, shown at e-value thresholds of $10^{-3}$, $10^{-1}$ and 10 (strict to lenient, left to right). All 13,035 ECFs are represented in each bar. The unmatched ('Neither') fraction decreased as the threshold was relaxed but did not disappear, remaining at 42% at an e-value of $\leqslant 0.1$ and 15% at the highly permissive threshold of $\leqslant 10$.

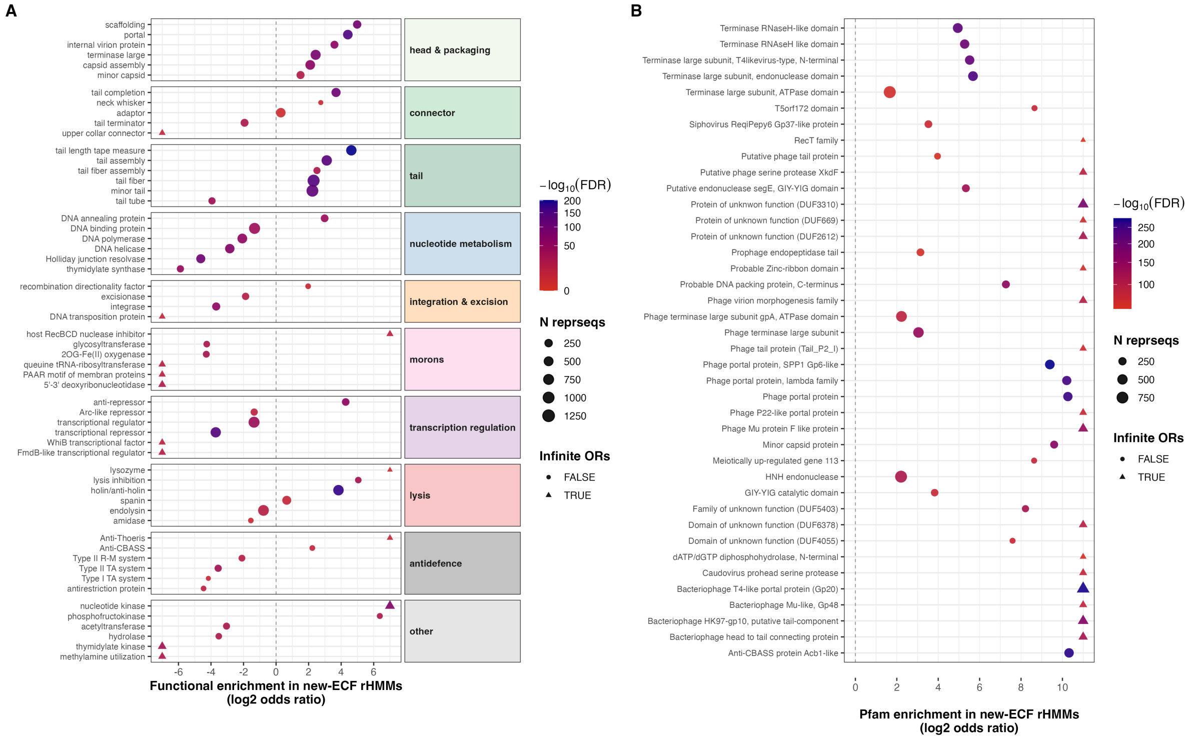


**Figure S11. Functional and Pfam enrichment among rHMMs carrying new ECFs.** New ECFs were defined as ECFs with no ECOD hit at an e-value of $\leqslant 0.1$, and an rHMM was treated as new-ECF-associated if it carried at least one such ECF. (A) PHROG functional enrichment. For each PHROG function, a two-sided Fisher's exact test compared the proportion of rHMMs carrying that function among new-ECF-associated rHMMs against all remaining annotated rHMMs; $p$-values were corrected using the Benjamini–Hochberg procedure. The six most significant functions per PHROG class are shown at FDR $\leqslant 0.05$, grouped by class in the row facets. Points give the $\log_2$ odds ratio, point size the total number of rHMMs carrying the function and colour the $-\log_{10}$ FDR. Infinite odds ratios are capped at $\pm 7$ and marked by a distinct point shape. The dashed line marks no enrichment. (B) Pfam enrichment, computed and displayed as in (A) but testing Pfam families instead of PHROG functions among the same set of rHMMs.

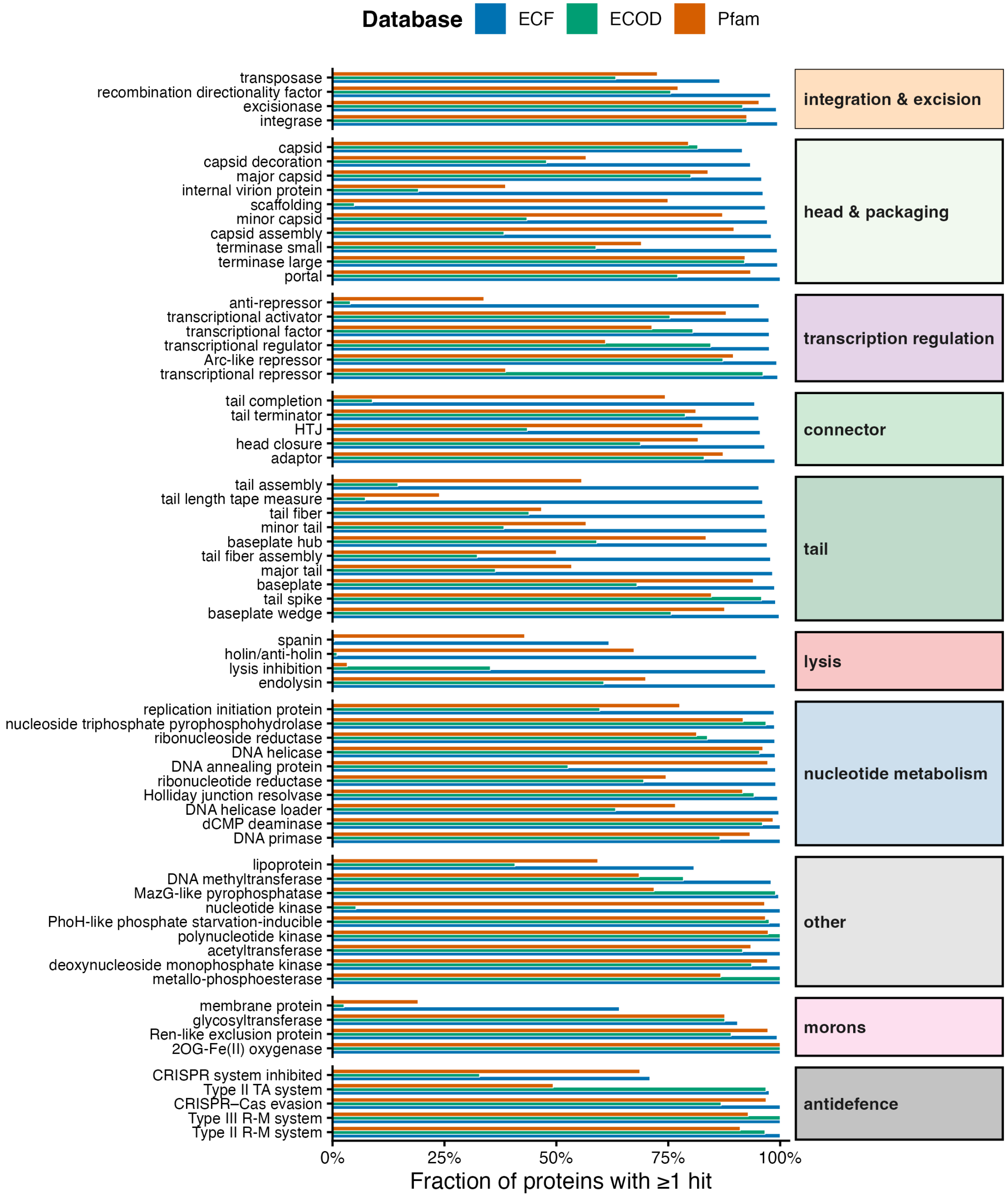


**Figure S12. Per-function annotation coverage of RefSeq phage proteins by ECF, ECOD, and Pfam.** Fraction of rHMMs with at least one hit from each database, shown for the top ten functions per PHROG functional class, ordered within each class by decreasing ECF coverage. Functional classes are indicated by coloured strip labels. Detection thresholds as in Figure 3.

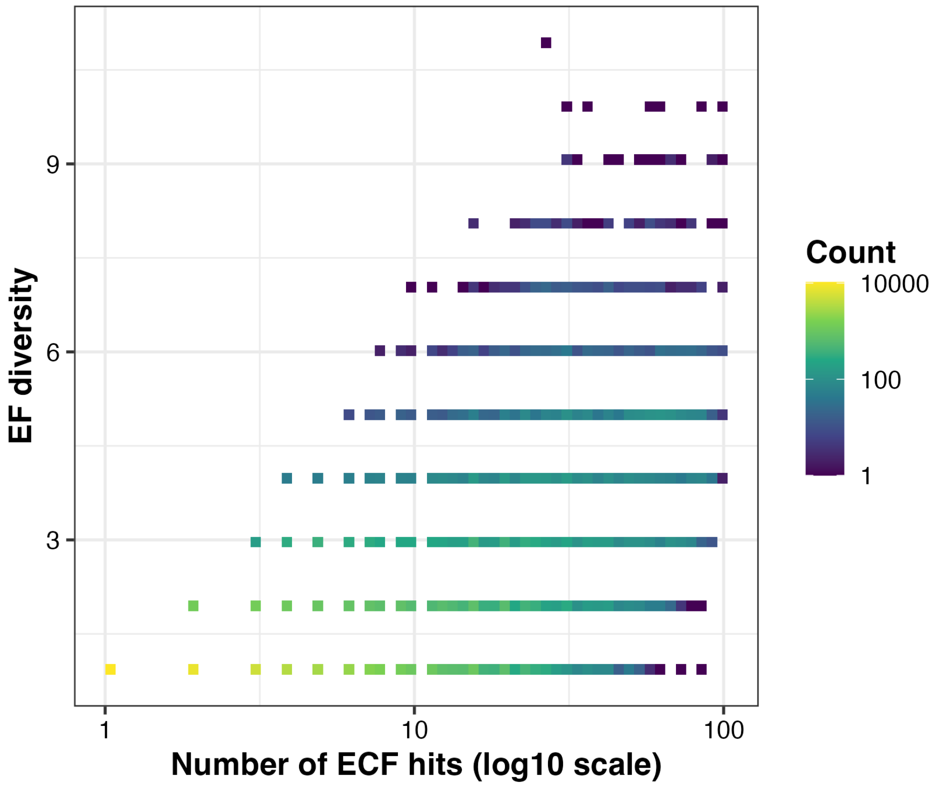


**Figure S13. EF diversity against detection depth in proteins shorter than 1,000 amino acids.** Binned density of the number of distinct EFs (`n.fams`) against the total number of ECF hits (`n.ecf.hits`, $\log_{10}$ scale), restricted to rHMMs shorter than 1,000 amino acids with non-zero ECF coverage. Bin counts are shown on a $\log_{10}$ viridis colour scale. The analysis repeats Figure 4A after removing exceptionally long proteins, which contribute the upper tail of the diversity distribution. The relationship between diversity and detection depth remained broad rather than deterministic, indicating that EF diversity was not simply a consequence of protein length.

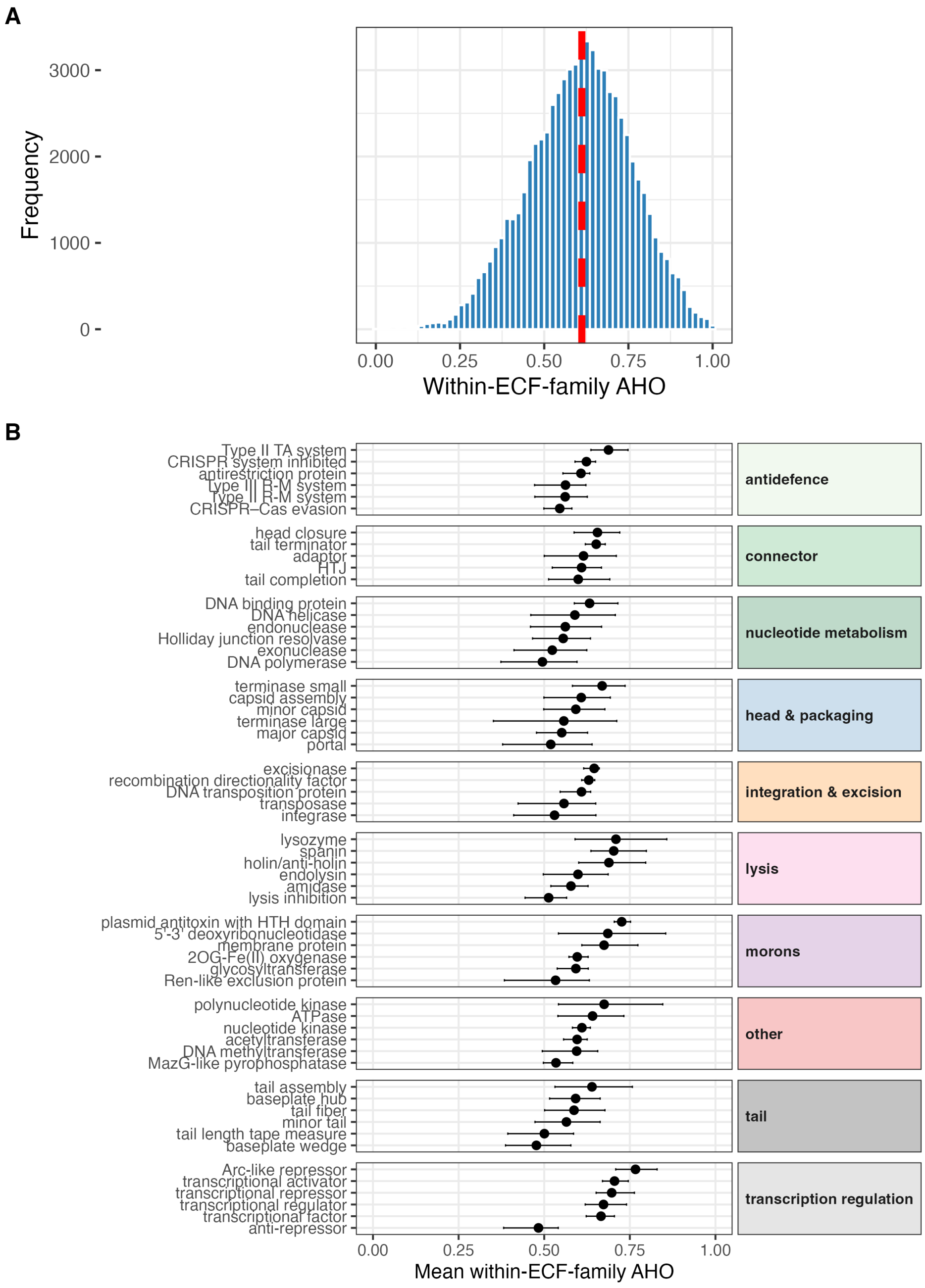


**Figure S14. Global distribution of within-ECF-family AHO.** (A) Histogram of mean within-ECF-family AHO across all rHMM–family pairs where the family contributed at least three ECF hits. The dashed red line marks the median. (B) Mean within-ECF-family AHO for the six most frequent functions per PHROG class. Points indicate the mean; horizontal bars span the interquartile range (25th–75th percentile). Only rHMM–family pairs where the family contributed at least three ECF hits were included. Functions are ordered by mean AHO within each class.

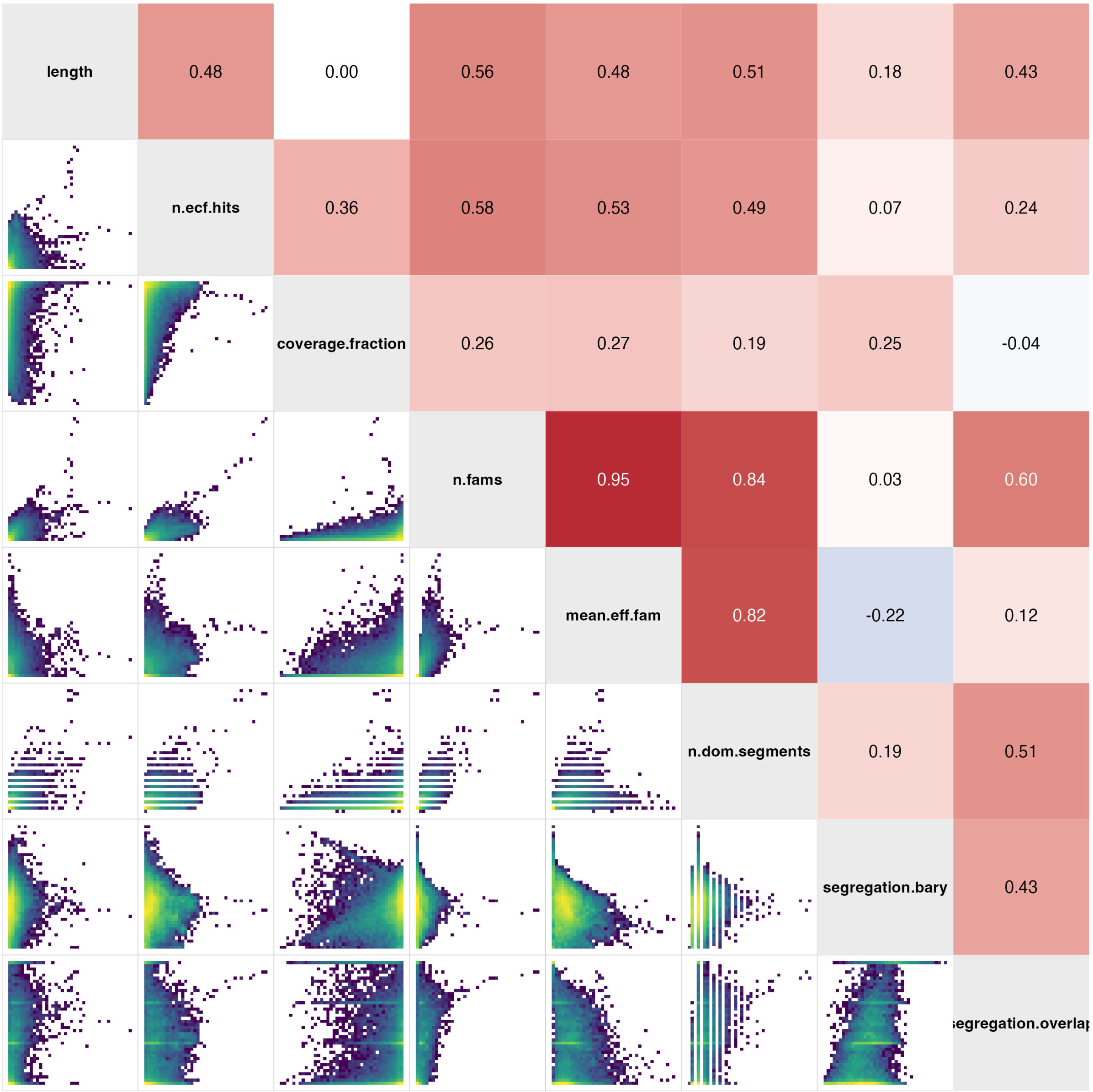


**Figure S15. Correlation structure of protein architecture metrics.** Pairwise Spearman correlations among eight per-rHMM architecture metrics: `coverage.fraction`, `length`, `mean.eff.fam`, `n.dom.segments`, `n.ecf.hits`, `n.fams`, `segregation.bary`, and `segregation.overlap`. Compositional complexity metrics (`n.fams`, `mean.eff.fam`, `n.dom.segments`) are strongly intercorrelated, whereas segregation metrics (`segregation.bary`, `segregation.overlap`) show weaker correlations with compositional complexity but moderate correlation with each other. This justifies the use of `n.fams` as a default measure of EF diversity and `segregation.bary` as the default measure of EF segregation.

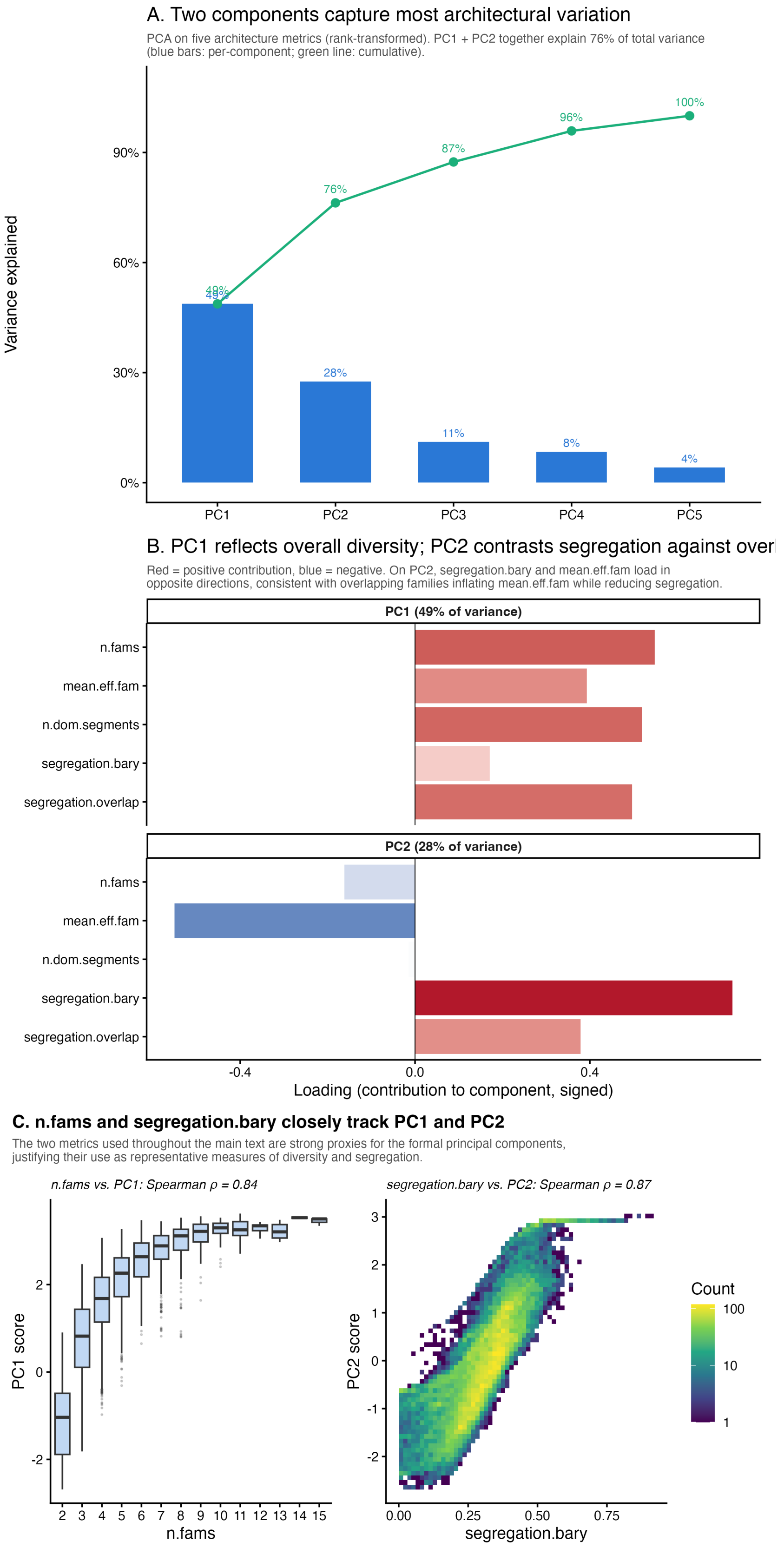
A. Two components capture most architectural variation
PCA on five architecture metrics (rank-transformed). PC1 + PC2 together explain 76% of total variance (blue bars: per-component; green line: cumulative).
Variance explained
0%
30%
60%
90%
49%
76%
87%
96%
100%
28%
11%
8%
4%
PC1
PC2
PC3
PC4
PC5
B. PC1 reflects overall diversity; PC2 contrasts segregation against overl
Red = positive contribution, blue = negative. On PC2, segregation.bary and mean.eff.fam load in opposite directions, consistent with overlapping families inflating mean.eff.fam while reducing segregation.
PC1 (49% of variance)
PC2 (28% of variance)
n.fams
mean.eff.fam
n.dom.segments
segregation.bary
segregation.overlap
-0.4
0.0
0.4
Loading (contribution to component, signed)
C. n.fams and segregation.bary closely track PC1 and PC2
The two metrics used throughout the main text are strong proxies for the formal principal components, justifying their use as representative measures of diversity and segregation.
n.fams vs. PC1: Spearman ρ = 0.84
segregation.bary vs. PC2: Spearman ρ = 0.87
PC1 score
n.fams
2 3 4 5 6 7 8 9 10 11 12 13 14 15
PC2 score
segregation.bary
0.00
0.25
0.50
0.75
Count
100
10
1

**Figure S16. Principal component analysis of per-rHMM architecture metrics.** Principal component analysis was performed on five rank-transformed architecture metrics (`n.fams`, `mean.eff.fam`, `n.dom.segments`, `segregation.bary` and `segregation.overlap`) across all rHMMs with complete values. Measures of detection depth and protein size (`n.ecf.hits`, `length`, `coverage.fraction`) were excluded, because including them merged depth with compositional complexity into the first component rather than leaving two interpretable architectural axes. (A) Scree plot. Bars give the variance explained by each component and the line gives the cumulative variance; the first two components together explained 76% of the total variance. (B) Loadings of each metric on PC1 and PC2, with facet labels giving the variance explained by each component. Red indicates a positive and blue a negative contribution. PC1 loaded consistently across the three compositional metrics, whereas on PC2 `segregation.bary` and `mean.eff.fam` loaded in opposite directions, consistent with overlapping families inflating the effective family count while reducing spatial separation. (C) Correspondence between the two metrics used in the main text and the formal components. Left, distribution of PC1 scores by `n.fams` (proteins with at most 15 families shown). Right, binned density of PC2 score against `segregation.bary`, with a $\log_{10}$ count colour scale. Spearman correlations are given above each panel.

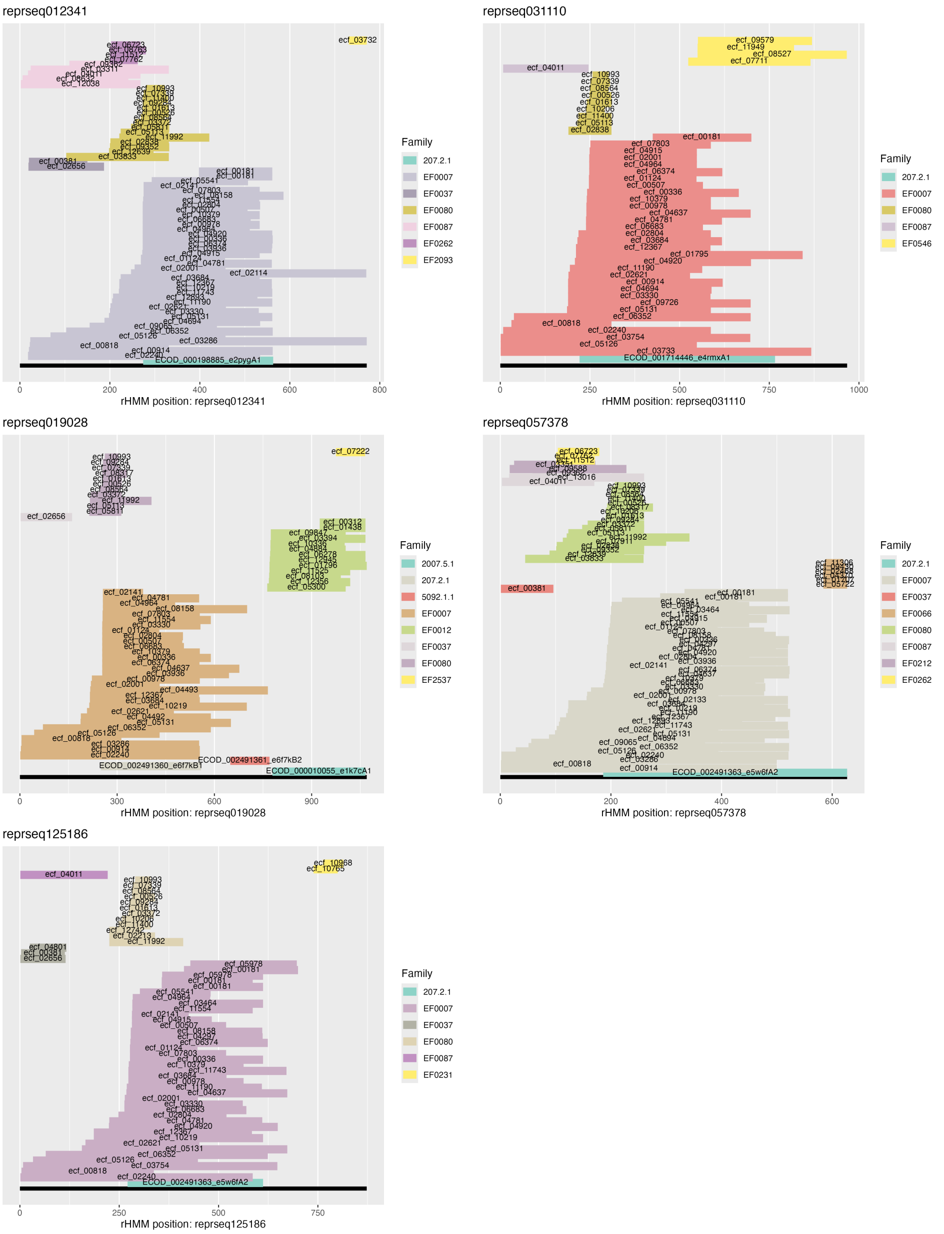


**Figure S17. Raw ECF and ECOD hit distributions across five depolymerase-type receptor-binding proteins shown in Figure 6.** All individual ECF hits detected in `reprseq012341`, `reprseq019028`, `reprseq031110`, `reprseq057378` and `reprseq125186`, plotted against position in the query rHMM. Each horizontal bar denotes a single ECF hit, labelled by its ECF identifier and coloured by ECF family (EF); hits are stacked vertically and ordered by start position. ECOD assignments are shown on the black backbone track below each panel, labelled by ECOD domain identifier and coloured by T-level using the same legend. Hits were retained at probability $\geqslant 95\%$, e-value $\leqslant 0.1$, minimum hit length $\geqslant 20$ aa and ECF coverage $\geqslant 70\%$. The consensus family blocks in Figure 6A were derived from these distributions by quartile-based boundary estimation; the unsmoothed hits are shown here to make hit density and boundary variability explicit.

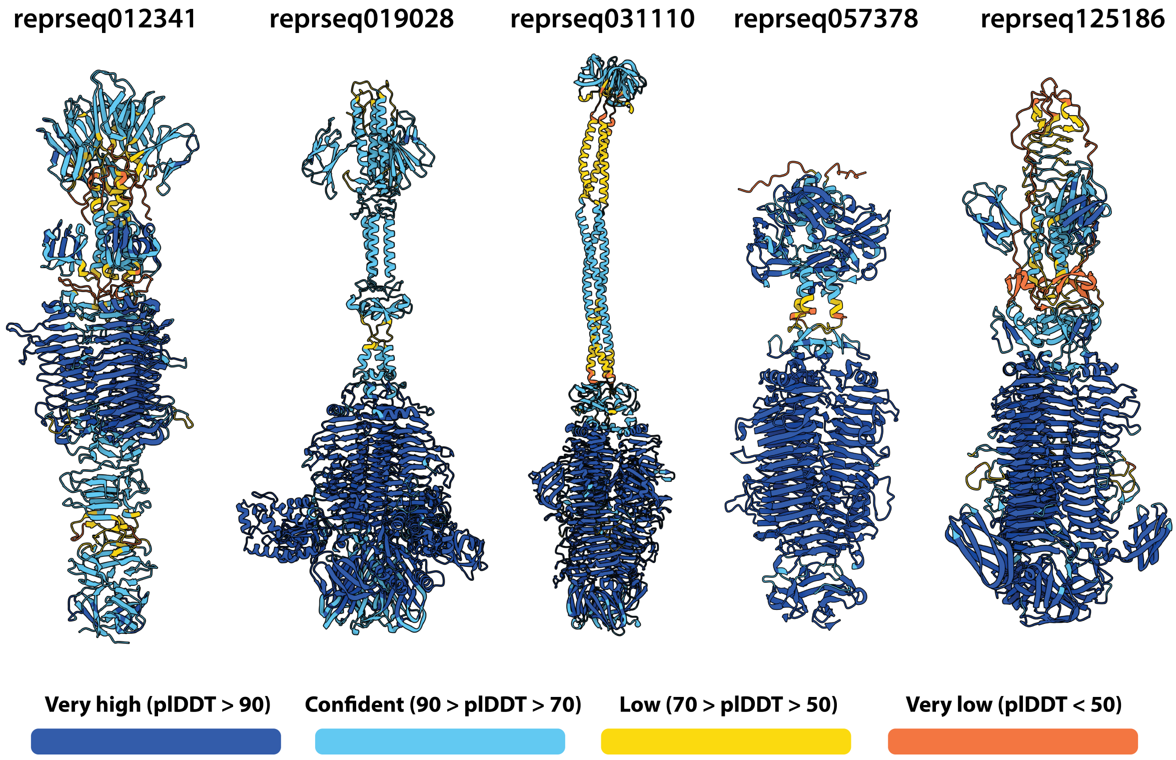


**Figure S18. AlphaFold3 models of five depolymerase-type receptor-binding proteins from Figure 6, coloured by predicted confidence.** Full-length homotrimeric models of `reprseq012341`, `reprseq019028`, `reprseq031110`, `reprseq057378` and `reprseq125186`, predicted with AlphaFold3[33] and coloured by per-residue pLDDT (dark blue, very high, pLDDT $>$ 90; light blue, confident, 70–90; yellow, low, 50–70; orange, very low, $<$ 50). Confidence was highest across the receptor binding domain (containing the single-stranded $\beta$-helix and the C-terminus), and fell in the extended N-terminal helical and coiled-coil segments, hence the model geometry in those regions should be interpreted accordingly. Figure 6B–D shows cropped regions of these models; complete structures are provided here for reference.

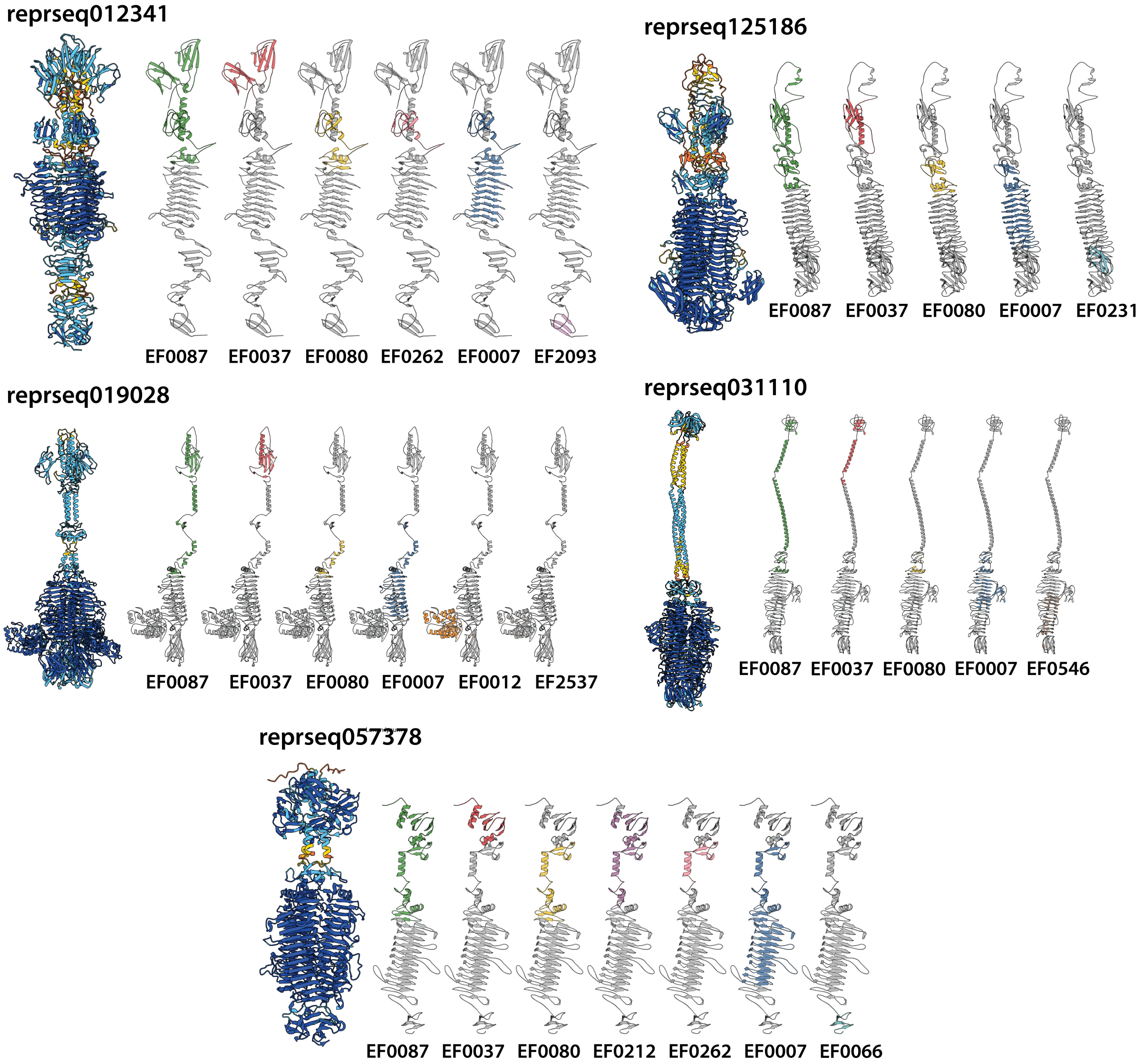


**Figure S19. ECF families mapped onto AlphaFold3 models of five depolymerase-type receptor-binding proteins.** For each protein, the complete homotrimeric model is shown on the left, coloured by pLDDT as in Supplementary Figure S18. To its right, chain A is shown once per detected ECF family, with residues assigned to that family coloured and all remaining residues in grey. Families are ordered from N- to C-terminus and labelled below each model. Assignments follow the consensus boundaries used in Figure 6A, derived from the raw hit distributions in Supplementary Figure S17. The regions shown in Figure 6B–D are drawn from these mappings.

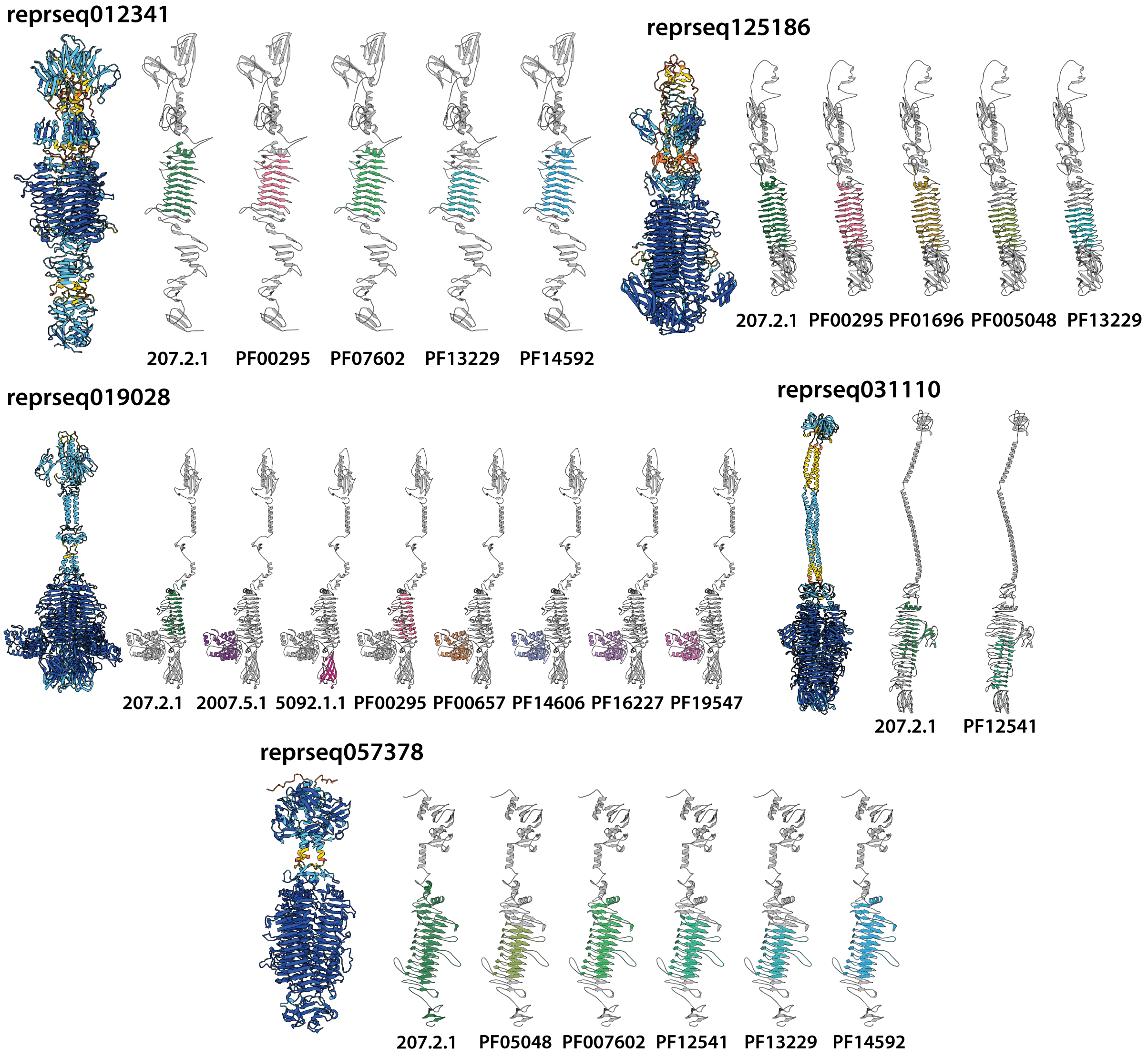


**Figure S20. ECOD and Pfam domains mapped onto AlphaFold3 models of five depolymerase-type receptor-binding proteins.** Layout as in Supplementary Figure S19: the complete model is shown on the left, followed by chain A with each detected ECOD or Pfam domain highlighted individually. Domains are ordered from N- to C-terminus and labelled below each model. Hits were detected under the thresholds applied throughout (probability $\geqslant 95\%$, e-value $\leqslant 0.1$, minimum hit length $\geqslant 20$ aa, domain coverage $\geqslant 70\%$). Comparison with Supplementary Figure S19 shows that several regions resolved into distinct ECF families received no assignment from either database.

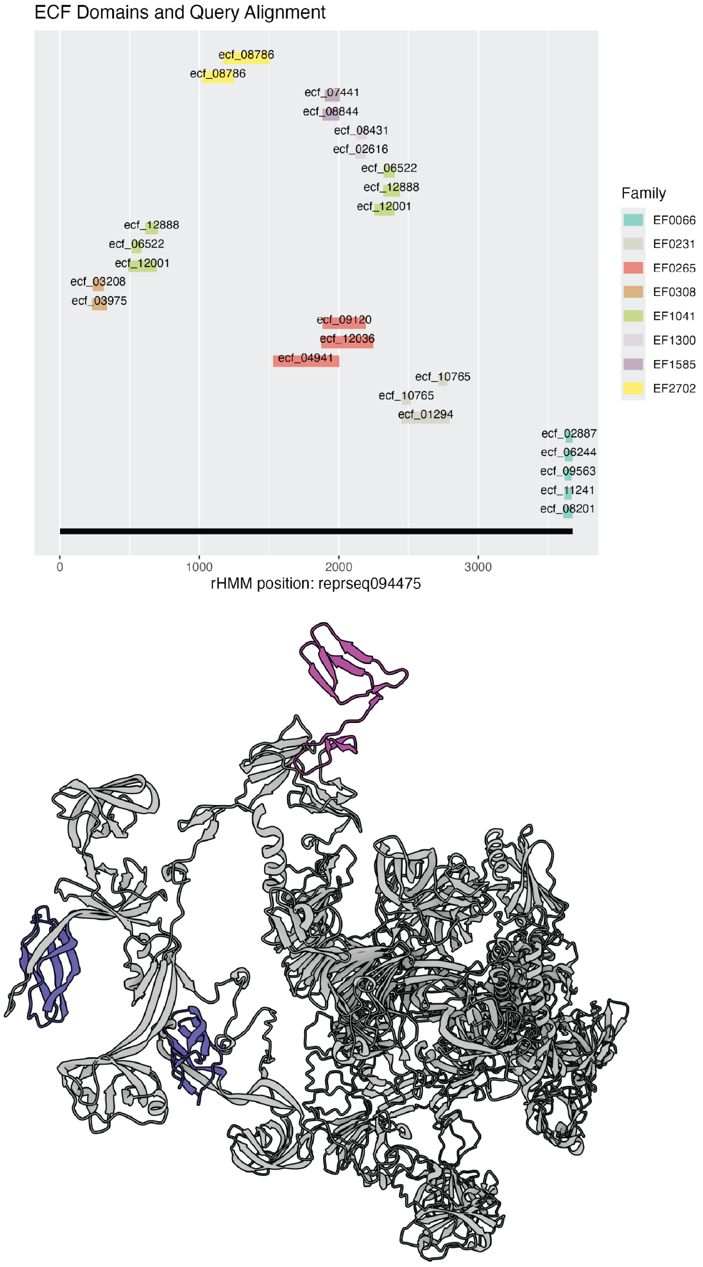


**Figure S21. ECF distribution and structural prediction of reprseq094475.** (Top) All individual ECF hits detected in `reprseq094475` plotted against position in the query rHMM, as shown and explained in Supplementary Figure S17. (Bottom) AlphaFold3 prediction of the protein sequence, highlighting the presence of EF0066 (magenta) and two occurrences of ecf_10765, assigned to EF0231 and found in `reprseq125186` (cf. Supplementary Figure S17).
